# The stem rust fungus *Puccinia graminis* f. sp. *tritici* induces centromeric small RNAs during late infection that direct genome-wide DNA methylation

**DOI:** 10.1101/469338

**Authors:** Jana Sperschneider, Ashley W. Jones, Jamila Nasim, Bo Xu, Silke Jacques, Narayana M. Upadhyaya, Rohit Mago, Melania Figueroa, Karam B. Singh, Eric A. Stone, Benjamin Schwessinger, Ming-Bo Wang, Jennifer M. Taylor, Peter N. Dodds

## Abstract

**Background:** Silencing of transposable elements (TEs) is essential for maintaining genome stability. Plants use small RNAs (sRNAs) to direct DNA methylation to TEs (RNA-directed DNA methylation; RdDM). Similar mechanisms of epigenetic silencing in the fungal kingdom have remained elusive.

**Results:** We use sRNA sequencing and methylation data to gain insight into epigenetics in the dikaryotic fungus *Puccinia graminis* f. sp. *tritici* (*Pgt*), which causes the devastating stem rust disease on wheat. We use Hi-C data to define the *Pgt* centromeres and show that they are repeat-rich regions (∼250 kb) that are highly diverse in sequence between haplotypes and, like in plants, are enriched for young TEs. DNA cytosine methylation is particularly active at centromeres but also associated with genome-wide control of young TE insertions. Strikingly, over 90% of *Pgt* sRNAs and several RNAi genes are differentially expressed during infection. *Pgt* induces waves of functionally diversified sRNAs during infection. The early wave sRNAs are predominantly 21 nts with a 5’ uracil derived from genes. In contrast, the late wave sRNAs are mainly 22 nt sRNAs with a 5’ adenine and are strongly induced from centromeric regions. TEs that overlap with late wave sRNAs are more likely to be methylated, both inside and outside the centromeres, and methylated TEs exhibit a silencing effect on nearby genes.

**Conclusions:** We conclude that rust fungi use an epigenetic silencing pathway that resembles RdDM in plants. The *Pgt* RNAi machinery and sRNAs are under tight temporal control throughout infection and might ensure genome stability during sporulation.

## Background

Epigenetic regulation controls transcription through formation of transcriptionally inactive chromatin (heterochromatin) and are mediated by interactions between small RNAs (sRNAs), DNA methylation and/or repressive histone modifications. In plants, sRNAs are predominantly in the size range of 20-24 nt and can be divided into two classes: 1) small interfering RNAs (siRNAs) processed by Dicer proteins from long double-stranded RNA (dsRNA), and 2) microRNAs (miRNAs) processed from stem-loop regions of single-stranded primary RNAs [1]. Both miRNAs and siRNAs are bound to argonaute (AGO) proteins to induce silencing of targets by base-pairing interactions and complementarity [2].

Heterochromatin plays both regulatory and structural roles. Heterochromatin regulates gene transcription, but also ensures proper chromosome segregation during cell division at centromeres and genome stability through regulation of transposable elements (TEs) [3]. Epigenetic silencing in repetitive genome regions is a key mechanism to prevent proliferation of TEs. In fungi and plants, DNA cytosine methylation (5-methylcytosine; 5mC) is found mainly in transposable elements and other repeated DNA [4, 5]. In plants, RNA-directed DNA methylation (RdDM) is the major sRNA-mediated epigenetic pathway and functions in maintaining genome stability through transposon control, pathogen defence and stress responses, intercellular communication and germ cell specification [6]. RdDM uses sRNAs to trigger DNA cytosine methylation at homologous DNA sequences in the genome [7]. These nuclear-localized heterochromatic sRNAs are the most abundant sRNAs in plants, predominantly 24 nucleotides (nts) in length, derived from intergenic or repetitive regions and associated with the argonaute 4 (AGO4) clade to regulate epigenetic silencing. Adenine is the most common 5′ base of AGO4-bound 24 nt sRNAs in *Arabidopsis* [8].

Unlike the extensively studied RdDM pathway in plants [9], the mechanisms of epigenetic silencing in the diverse fungal kingdom have remained elusive [10]. The RNAi machinery of the fission yeast *Schizosaccharomyces pombe* and the filamentous fungus *Neurospora crassa* are thus far the best-studied non-pathogenic model species [11]. In fission yeast, RNAi components participate in heterochromatin formation through histone H3K9 modifications at centromeres, the mating type interval and the subtelomeric regions [12, 13]. DNA cytosine methylation is absent in the model yeasts *S. pombe* and *S. cerevisiae* [14]. In *Neurospora crassa*, RNAi components are involved in quelling and meiotic silencing by unpaired DNA (MSUD), but not in heterochromatin formation. Quelling is an RNAi-related gene-silencing mechanism in *Neurospora* that is induced by repetitive transgenic sequences and occurs in the vegetative growth stage to control transposons [15]. Outside the model fungal species, very little is known about the interplay between sRNAs and epigenetic silencing, particularly in highly repetitive fungal pathogen genomes that need to inactivate TEs. Unlike plants, fungi lack canonical gene-body methylation but in line with plants, 5mC is abundant in repetitive DNA and transposons across fungal species [4]. How RNAi contributes to epigenetic silencing in other fungi has thus far largely remained unexplored. In the ascomycete *Magnaporthe oryzae*, a plant pathogen, 18-23 nt sRNAs are produced from repetitive elements and are implicated in TE regulation in vegetative tissue, whereas 28-35 nt sRNAs mapping to transfer RNA (tRNA) loci are enriched in the appressoria [16]. However, a correlation between sRNAs and epigenetic silencing has not been shown in *M. oryzae*. In the white-rot basidiomycete fungus *Pleurotus ostreatus*, TE silencing is associated with 21 nt sRNAs and DNA methylation [17]. RNAi has been suggested as a key determinant of longer centromeres in the human fungal pathogen *Cryptococcus* and as a suppressor of centromeric retrotransposons to ensure genome stability [18].

The basidiomycete fungus *Puccinia graminis* f. sp. *tritici* (*Pgt*) is a plant pathogen that causes wheat stem rust disease, resulting in devastating crop losses [19]. *Pgt* is a dikaryotic fungus that contains two distinct haploid nuclei. During the asexual infection phase on a cereal host, *Pgt* produces single-celled dikaryotic urediniospores that germinate on the leaf surface [20, 21]. Subsequently, appressoria form and penetration occurs through stomata with subsequent development of specialized infection structures called haustoria by around 2 days. Haustoria enable nutrient uptake as well as the delivery of secreted pathogen proteins called effectors into the host plant cell [22]. The start of urediniospore production occurs at approximately 6-7 days post infection (dpi) and urediniospore pustules typically erupt through the leaf or stem surface (sporulation) after 8–10 dpi [20]. In the poplar rusts, intense cell division activity has been observed in the sporulation area [23].

Chromosome-scale genome assemblies offer the opportunity to investigate the structural organisation of the genome including localization of centromeres, transposable elements (TEs), DNA methylation, sRNAs and how this links to their function. Recently, the chromosome-scale assembly of *Pgt* 21-0 has become available [24]. This assembly is fully phased with 18 chromosome pseudomolecules for each of the two haplotypes derived from the two haploid nuclei. Whilst substantial time-course transcriptomic resources have been generated for *Pgt* [25-27], how it utilizes RNAi and epigenetic silencing during the infection cycle has thus far been unknown. Here, we bring together Hi-C data, methylation data and sRNA/transcriptome sequencing data to uncover fundamental characteristics of the stem rust RNAi machinery, DNA methylation and the first-time characterization of rust centromeres.

## Results

### *Pgt* centromeres are highly repetitive sequences with little sequence conservation between haplotypes

We used chromatin conformation capture assay data (Hi-C) from *Pgt* 21-0 [24] to pinpoint the location of the *Pgt* centromeres, the first-time characterization of rust fungi centromeres. Fungal centromeres give rise to a distinct outwards-spreading shape in a Hi-C contact map [28], as seen in the contact maps for individual chromosomes (Supplementary Figures S1 and S2). Because the centromeres of each chromosome tend to cluster in the three-dimensional space of the nucleus, this region also shows physical association between chromosomes visible as distinct cross-shapes in the whole haplotype Hi-C contact map (Figure 1A). We selected the midpoint of the outwards-spreading shape in the Hi-C contact maps of each chromosome as the putative centre of each centromere. For example, *Pgt* chromosome 1A has a suggested centromere centre around position 2.36 MB and the surrounding region shares strong Hi-C links with the putative centromeres on other chromosomes (Figure 1A and Supplementary Figure S3). To add further support to the centromeric regions and their lengths, we plotted the density of expressed genes, RNA-seq transcription levels at late infection and in germinated spores as well as the coverage of repetitive elements on the chromosomes. This shows that the regions around the putative centromere centres indicated by the Hi-C data are transcriptionally silent, gene-poor and repeat-rich regions (Figures 1B and 2, Supplementary Figure S4). We selected putative centromere boundaries by inspecting the lengths of these transcriptionally silent, gene-poor regions for each chromosome (Figure 2). Centromeres appear to span between 100 kb to 340 kb (253 kb on average), with only a slight decrease in GC content for most chromosomes compared to the rest of the chromosome (Table 1).

**Table 1:** Genomic coordinates, lengths and GC content of the centromeric regions for each *Pgt* chromosome of the A and B haplotypes.

| Chromosome | Centromeric region | Centromere length | GC content<br>(centromere/non-centromere) |
| --- | --- | --- | --- |
| 1A | 2.18 – 2.45 MB | 270 kb | 40.6% / 43.7% |
| 1B | 2.44 – 2.72 MB | 280 kb | 41.9% / 43.6% |
| 2A | 1.66 – 1.93 MB | 270 kb | 40.9% / 43.8% |
| 2B | 1.56 – 1.83 MB | 270 kb | 41.8% / 43.7% |
| 3A | 1.87 – 2.09 MB | 220 kb | 41.9% / 43.8% |
| 3B | 1.92 – 2.26 MB | 340 kb | 43.4% / 43.3% |
| 4A | 3.06 – 3.34 MB | 280 kb | 42.1% / 43.7% |
| 4B | 3.33 – 3.64 MB | 310 kb | 42.6% / 43.5% |
| 5A | 4.38 – 4.65 MB | 270 kb | 42.3% / 43.8% |
| 5B | 6.01 – 6.23 MB | 220 kb | 42.4% / 43.9% |
| 6A | 1.30 – 1.62 MB | 320 kb | 43.3% / 43.5% |
| 6B | 1.15 – 1.49 MB | 340 kb | 42.8% / 43.2% |
| 7A | 1.74 – 2.04 MB | 300 kb | 41.9% / 43.6% |
| 7B | 1.94 – 2.18 MB | 240 kb | 41.1% / 43.4% |
| 8A | 1.24 – 1.52 MB | 280 kb | 41.6% / 43.6% |
| 8B | 1.28 – 1.55 MB | 270 kb | 42.5% / 43.9% |
| 9A | 1.93 – 2.16 MB | 230 kb | 41.9% / 44.0% |
| 9B | 2.12 – 2.36 MB | 240 kb | 41.9% / 43.7% |
| 10A | 1.37 – 1.66 MB | 290 kb | 42.8% / 43.7% |
| 10B | 1.98 – 2.25 MB | 270 kb | 42.9% / 43.4% |
| 11A | 3.5 – 3.6 MB | 100 kb | 40.5% / 43.8% |
| 11B | 3.82 – 3.93 MB | 110 kb | 40.5% / 43.7% |
| 12A | 2.85 – 3.13 MB | 280 kb | 41.8% / 43.6% |
| 12B | 2.82 – 3.08 MB | 260 kb | 40.8% / 43.5% |
| 13A | 1.9 – 2.14 MB | 240 kb | 41.7% / 43.5% |
| 13B | 2.06 – 2.28 MB | 220 kb | 41.4% / 43.6% |
| 14A | 2.13 – 2.44 MB | 310 kb | 42.0% / 43.6% |
| 14B | 2.25 – 2.48 MB | 230 kb | 42.8% / 43.4% |
| 15A | 2.62 – 2.78 MB | 160 kb | 42.5% / 43.6% |
| 15B | 2.52 – 2.81 MB | 290 kb | 41.3% / 43.5% |
| 16A | 2.3 – 2.54 MB | 240 kb | 41.0% / 43.9% |
| 16B | 2.26 – 2.55 MB | 290 kb | 42.5% / 43.7% |
| 17A | 2.35 – 2.52 MB | 170 kb | 42.5% / 43.7% |
| 17B | 2.18 – 2.45 MB | 270 kb | 41.5% / 43.7% |
| 18A | 1.79 – 2.02 MB | 230 kb | 43.3% / 43.7% |
| 18B | 2.02 – 2.22 MB | 200 kb | 42.3% / 43.4% |

**Figure 1:**
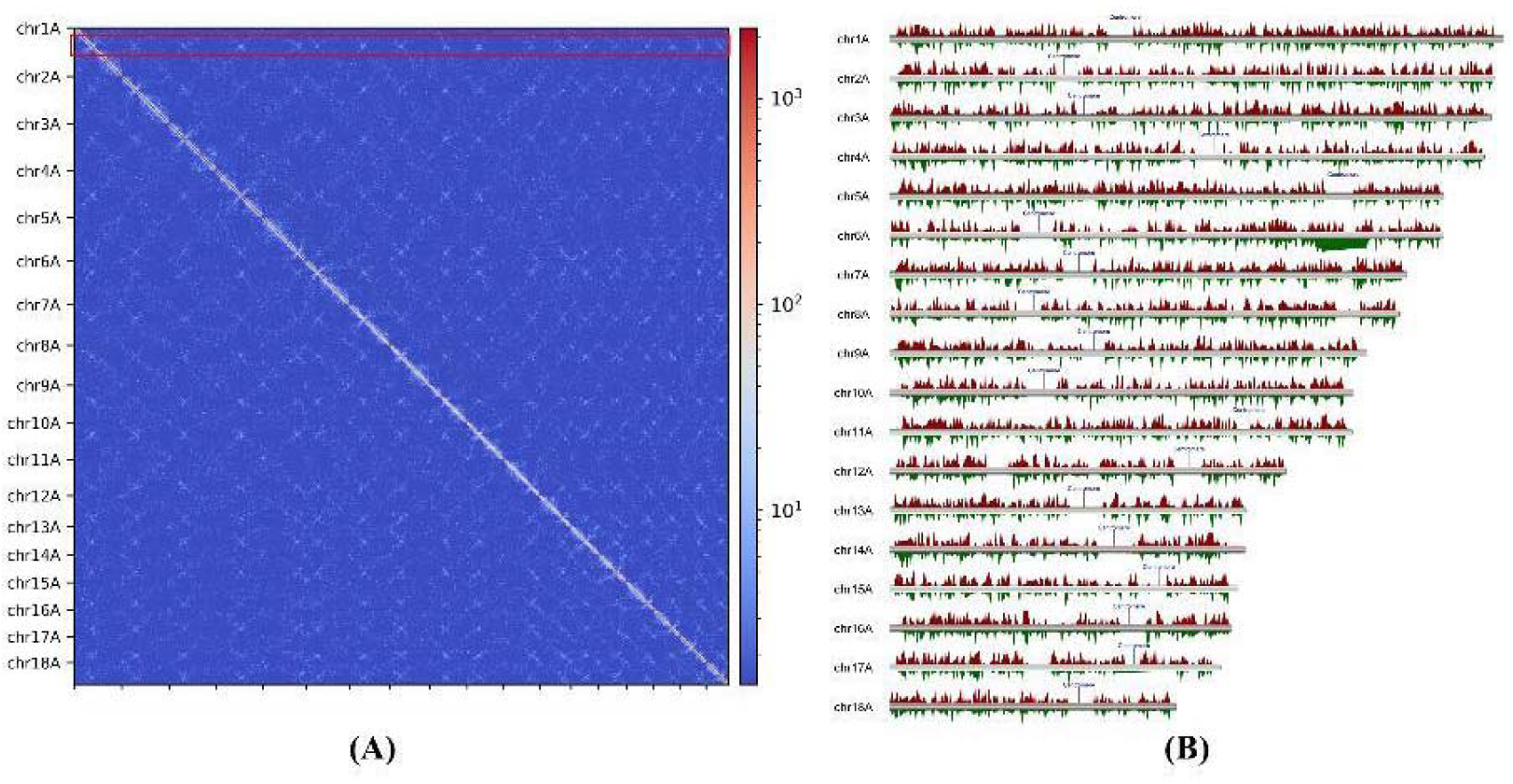
Hi-C contact map shows the location of the *Pgt* centromeres. **(A)** A Hi-C contact map of the 18 chromosomes in haplotype A shows the position of the centromeres as cross-like shapes, highlighted with a red rectangle. **(B)** The positions of the centromeres in haplotype A as indicated by the Hi-C contact map are in transcriptionally silent genomic regions. Reads per million (RPM) for the late infection (7 dpi) and germinated spores RNAseq samples are shown in red and green, respectively (10 kb windows, RPM from 0-100 are shown for clarity).

**Figure 2:**
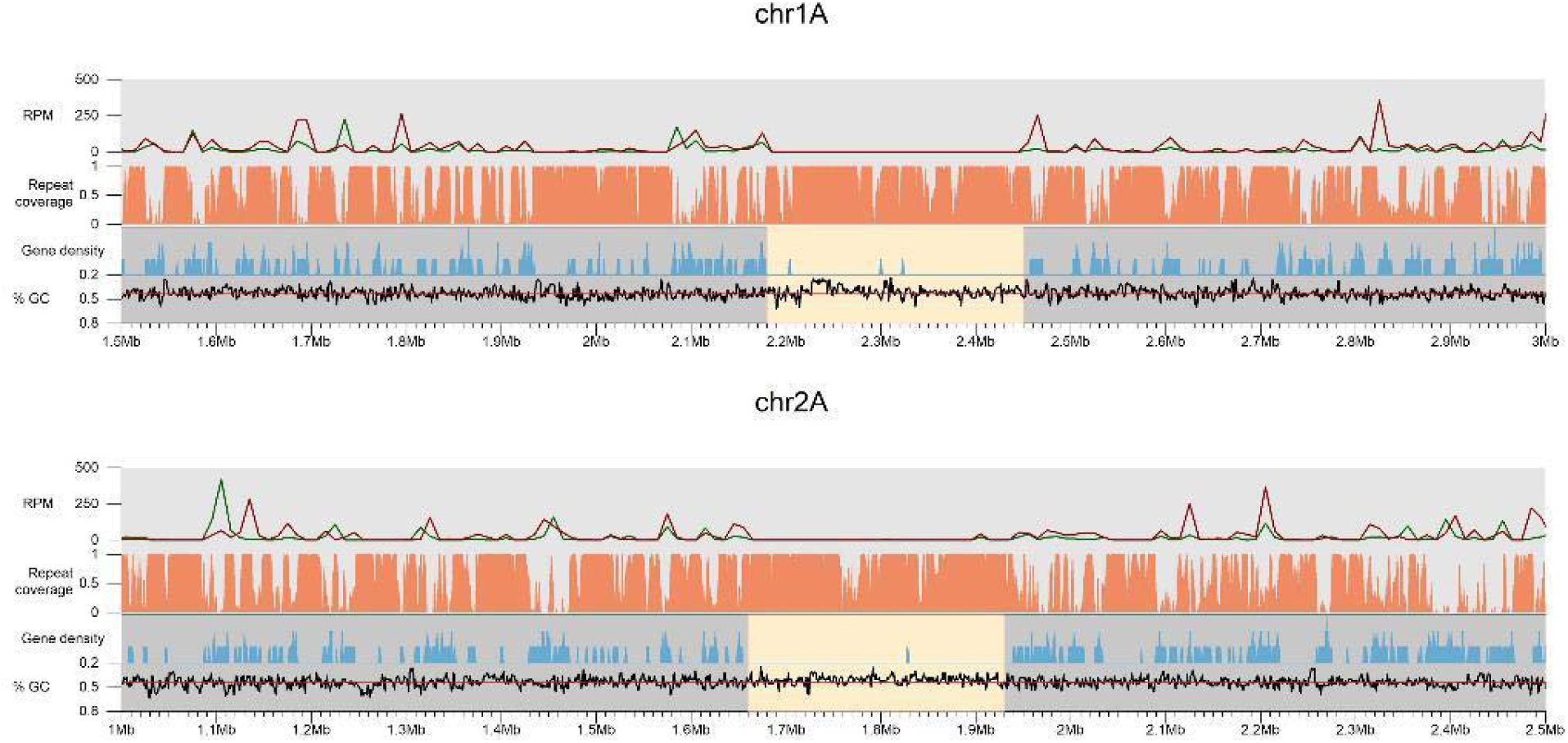
*Pgt* centromeric regions for two selected chromosomes. Karyoplots of the centromeric regions of *Pgt* chromosomes 1A and 2A. The density of expressed genes and the coverage of repetitive elements are shown as well as the GC content (1 kb windows). Reads per million (RPM) for the late infection (7 dpi) and germinated spores RNAseq samples are shown as red and green lines, respectively (10 kb windows). Centromeric regions are highlighted with yellow boxes.

We then aligned the two haplotype chromosomes. Interestingly, some chromosomes share regions of macro-synteny (conserved regions > 20 kb) in their centromeres whereas others do not. For example, chromosomes 1A and 1B show no to very low sequence identity in the centromeric region, as opposed to the remainder of the chromosome (Figure 3A). In contrast, chromosomes 2A and 2B share macro-synteny in the centromeric region (Figure 3B). We then used pairwise *k*-mer distance estimation to compare centromeric regions and non-centromeric regions for all chromosomes. Clustering analysis showed a large distance between the centromeric regions and non-centromeric regions (Figure 3C). For the non-centromeric regions, the two homologous chromosomes all grouped into closely related pairs with similar distances for all 18 chromosome pairs. In contrast, while most (14/18) centromeric regions grouped by chromosome pairs, the difference between them varied, with some very closely related (e.g. on chromosomes 2A and 2B) and others quite divergent (e.g. chromosomes 1A and 1B). Others showed unexpected groupings. For example, the centromeres of chromosomes 18A and 15B are more closely related to each other than to the centromeres of the corresponding chromosomes 18B and 15A (Figure 3C). Overall, the centromeric regions of *Pgt* are highly variable and unexpectedly, most of them are also highly divergent between haplotypes.

**Figure 3:**
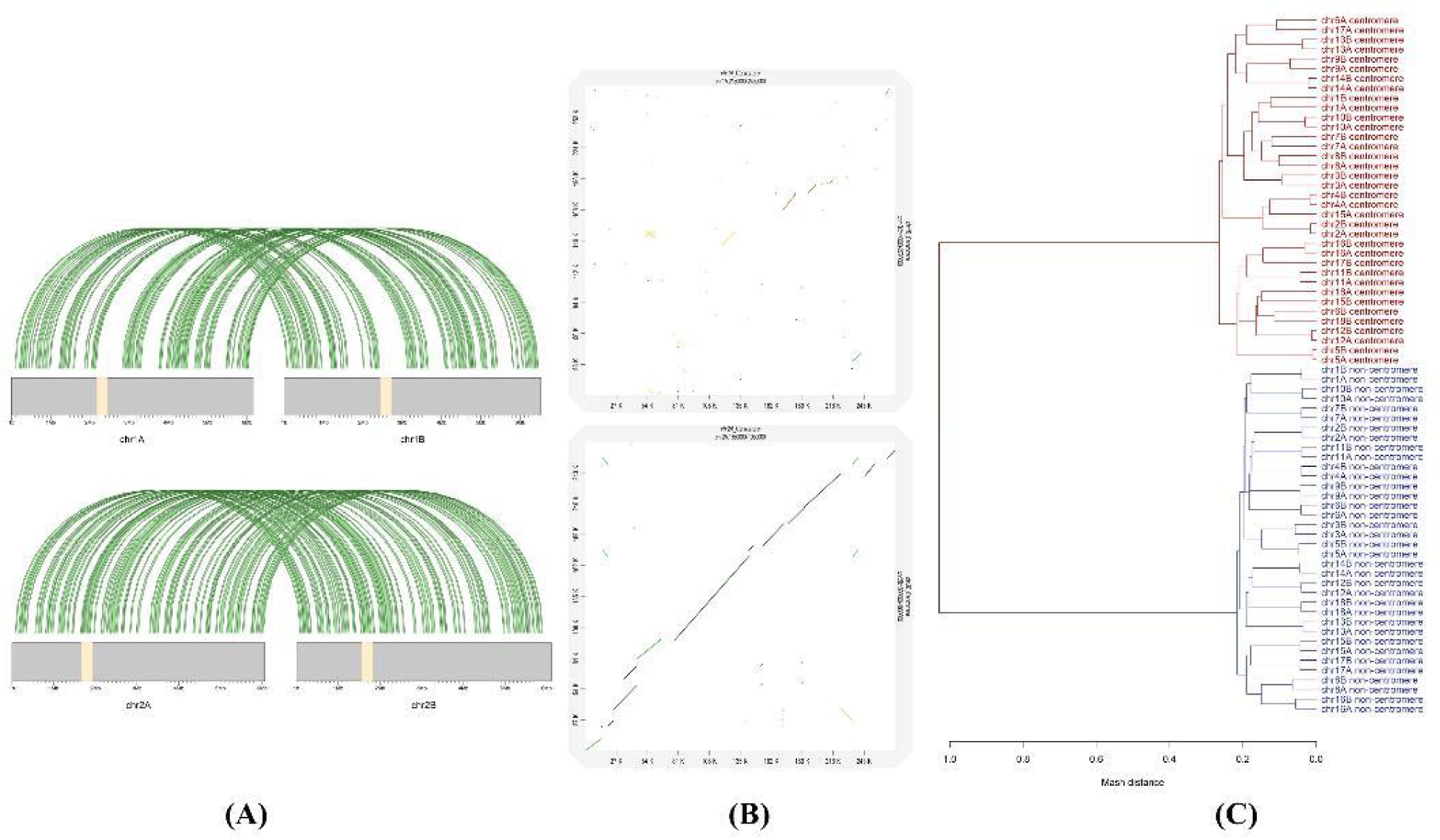
Synteny and sequence similarity of the *Pgt* centromeres. **(A)** Regions of macrosynteny (> 20 kbp) between the haplotypes of chromosomes 1 and 2. For chromosome 1, the centromeric regions show no conservation whereas on chromosome 2 the centromeric regions are conserved as confirmed by **(B)** genomic dot plot alignments of the centromeric regions. **(C)** Clustering of *k*-mer distance estimations between centromeric and non-centromeric regions. The non-centromeric regions cluster as expected according to haplotypes. In contrast, the centromeric regions are highly divergent, even between haplotypes.

### Young transposable elements accumulate in the highly repetitive *Pgt* centromeric regions

We then assessed the repetitive element coverage of the *Pgt* chromosomes and their centromeres. All 36 (2*18) *Pgt* centromere regions have higher coverage of repetitive elements than the non-centromeric regions (Figure 4A). Repetitive elements cover 75%-96% of the bases in the *Pgt* centromeres compared to only 52%-62% of the non-centromeric regions on the chromosomes. The repeat types with the highest sequence coverage in the centromeric regions vary considerably between the chromosomes. Most centromeres are enriched for LTR Gypsy retrotransposons (17%-56%) compared to non-centromeric regions (11-17%), although this is also the most abundant repeat family outside the centromeres (Figure 4B). However, several centromeres show a high coverage of repeat families that are of low abundance outside the centromeric regions. For example, DNA transposons of the superfamily CACTA are highly abundant in the centromere of chromosome 17A (33% coverage), while the centromere of chromosome 11B is enriched for LTR Copia retrotransposons (35% coverage). Again, these patterns are not always shared between centromeres within a chromosome pair.

**Figure 4:**
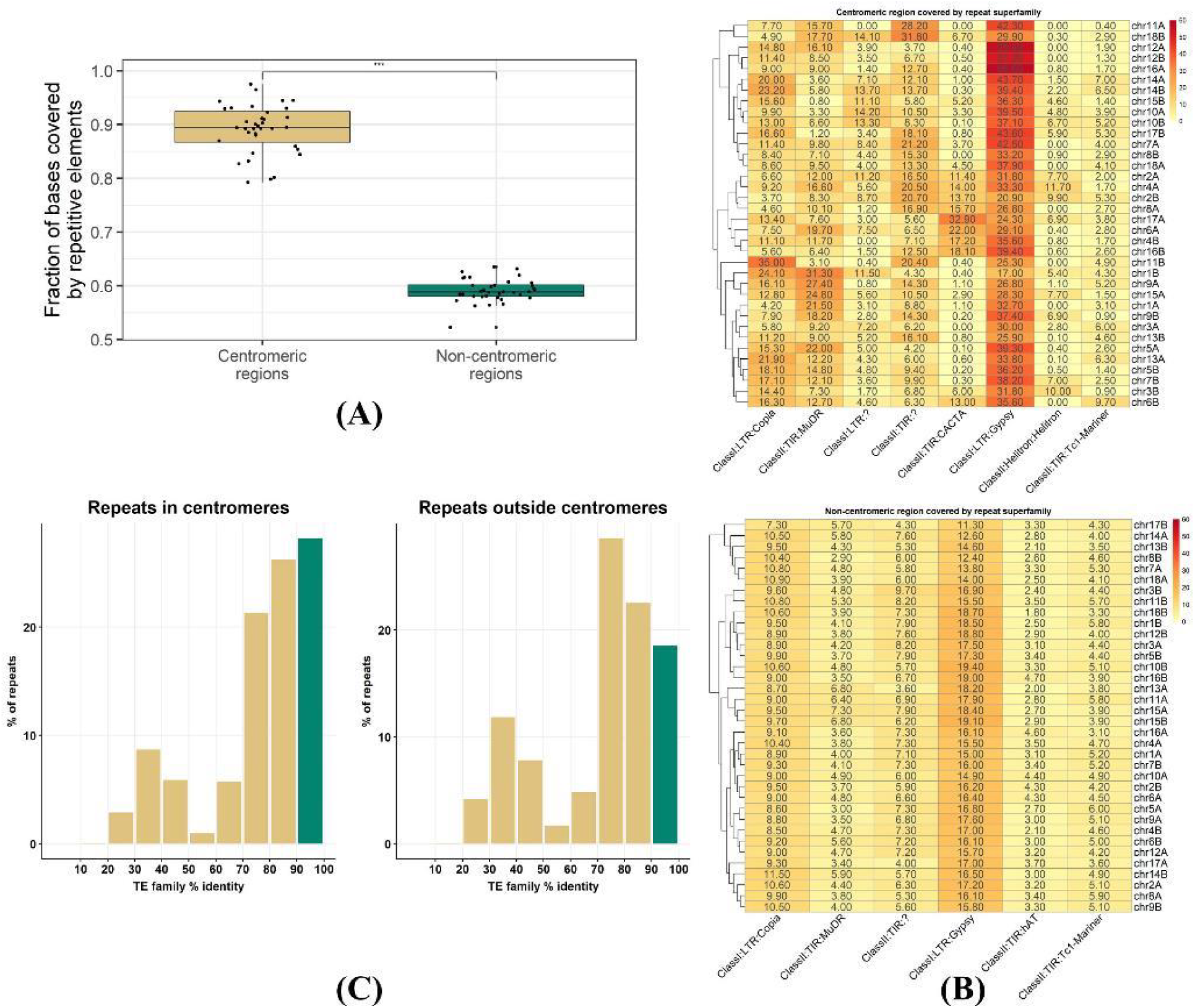
Properties of repetitive elements in the centromeres. **(A)** The repetitive element coverage of centromeric regions is significantly higher than the repetitive element coverage of non-centromeric regions for all the *Pgt* chromosomes. **(B)** Percent of bases that are covered by repetitive elements of a particular class. The centromeric regions vary considerably in the types of repeats they harbour, even between haplotypes. **(C)** The centromeres have a large proportion of young transposable element (TE) insertions compared to the non-centromeric regions.

To determine whether the age of TEs affects their distribution, we used the nucleotide identity of each TE to the consensus sequence of the family as a proxy for the relative age of TE insertion. Most TEs in the *Pgt* genome have >70% identity to the consensus, however the centromeres are enriched for young TEs (defined as having > 90% identity, Figure 4C). In the centromeres, 28.3% of repeats with family identity information are young TEs compared to 18.8% outside the centromeres. Taken together, the centromeres are highly repetitive regions in the *Pgt* genome that are enriched for young TEs, similarly to *Arabidopsis* where the majority of young repeats are found in pericentromeric domains [29].

### *Pgt* centromeres are heavily 5mC-methylated at genomic CG sites

We used Nanopore sequencing to detect DNA methylation in genomic DNA of *Pgt* during two distinct infection stages: 1) germination of spores and 2) late infection stage of wheat when sporulation starts (7 dpi). The Nanopore signal distinguishes 5-methylcytosine (5mC) from unmethylated cytosine and N6-methyladenine (6mA) from unmethylated adenine [30]. Methylated sites were defined as nucleotide positions where more than 50% of sequence reads showed the presence of a modified residue. We found that the occurrence of methylated cytosine residues was substantially higher in CG dinucleotide and CCG trinucleotides than in other di- and trinucleotide contexts, similar to cytosine methylation patterns in plants. The proportion of both CG and CCG methylation sites was significantly higher in centromeric regions (18% and 7.5%) than in non-centromeric regions (7.5% and 3.6%). Levels of 6mA methylation were low both inside and outside the centromeres with no substantial difference between dinucleotide contexts (Figure 5A). The frequency of methylation at methylated CG dinucleotide sites (i.e. % of reads from a site that show base modifications) are also higher for sites that occur in the centromeres than for those outside the centromeres, but very similar between germinated spores and late infection (Figure 5B). Taken together, *Pgt* has a strong preference for genomic CG (CpG) methylation and centromeres are heavily CG-methylated genomic regions.

**Figure 5:**
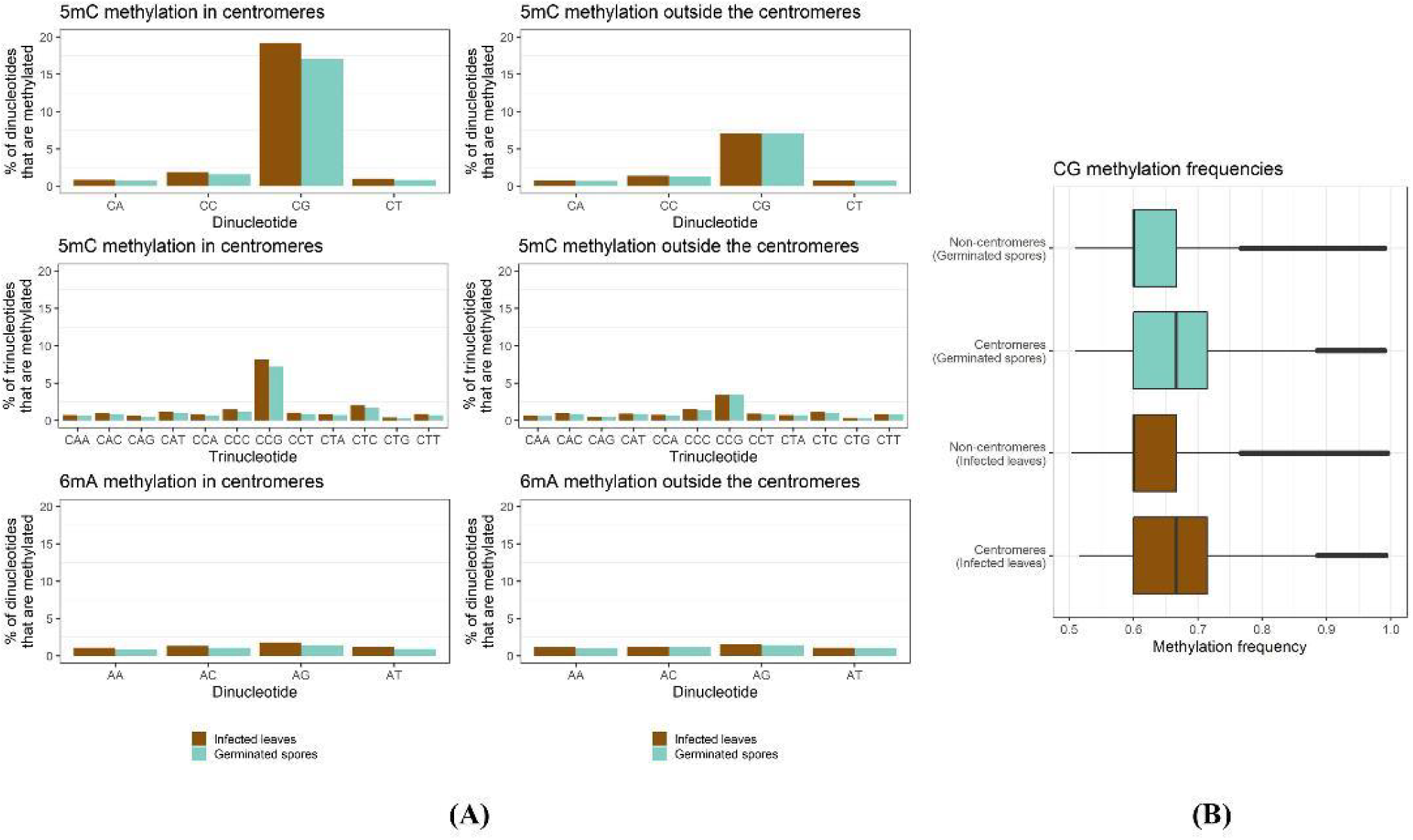
Methylation site preferences in *Pgt*. **(A) Proportions of *Pgt* dinucleotides and trinucleotides that are methylated in the centromeres and outside the centromeres.** CG dinucleotides are highly enriched for 5mC methylation. For the trinucleotides, CCG is enriched for 5mC methylation. We observed very low levels of 6mA methylation. The centromeres are heavily methylated compared to the non-centromeric regions. Slightly higher levels of 5mC methylation are seen in infected leaves compared to germinated spores in centromeres. **(B) Box plots showing methylation frequency distribution of CG methylation sites.** Centromeres show higher methylation frequencies than non-centromeric regions.

### CG methylation is associated with silencing of young TE insertions both inside and outside of centromeres

CG methylation is strongly associated with repetitive regions in both germinated spores and late infection, with 89.3% and 88.3% of methylation sites overlapping with TEs, respectively. 12.2% of all *Pgt* TEs are methylated in germinated spores and 12.2% in late infection. 47% of methylated TEs show methylation in both conditions. A higher percentage of TEs in centromeric regions are methylated (21.4% and 17.2% in infected leaves and germinated spores, respectively) than TEs in non-centromeric regions (11.8% and 12% in infected leaves and germinated spores, respectively). We did not observe significant differences in TE family distributions for TEs that are methylated only in germinated spores or only in infected leaves (data not shown).

CG methylation is strongly associated with young TEs (> 90% identity), not only inside the centromeric regions but even more so outside the centromeres. 21.2% and 25.1% of young TEs in the centromeres are methylated in germinates spores and in late infection, respectively (Figure 6). Whilst the centromeres are highly methylated genomic regions and preferentially harbour young TE elements (Figure 4C), young TEs that occur outside the centromeres are also heavily methylated. 23.6% of young TEs outside the centromeres are methylated both in germinated spores and in infected leaves (Figure 6). This suggests that *Pgt* employs a mechanism that maintains silencing of young TEs not only in the centromeres but also through targeting of their homologous sequences outside the centromeres. We hypothesized that *Pgt* might employ sRNAs to direct DNA methylation to young TEs outside the centromeres.

**Figure 6:**
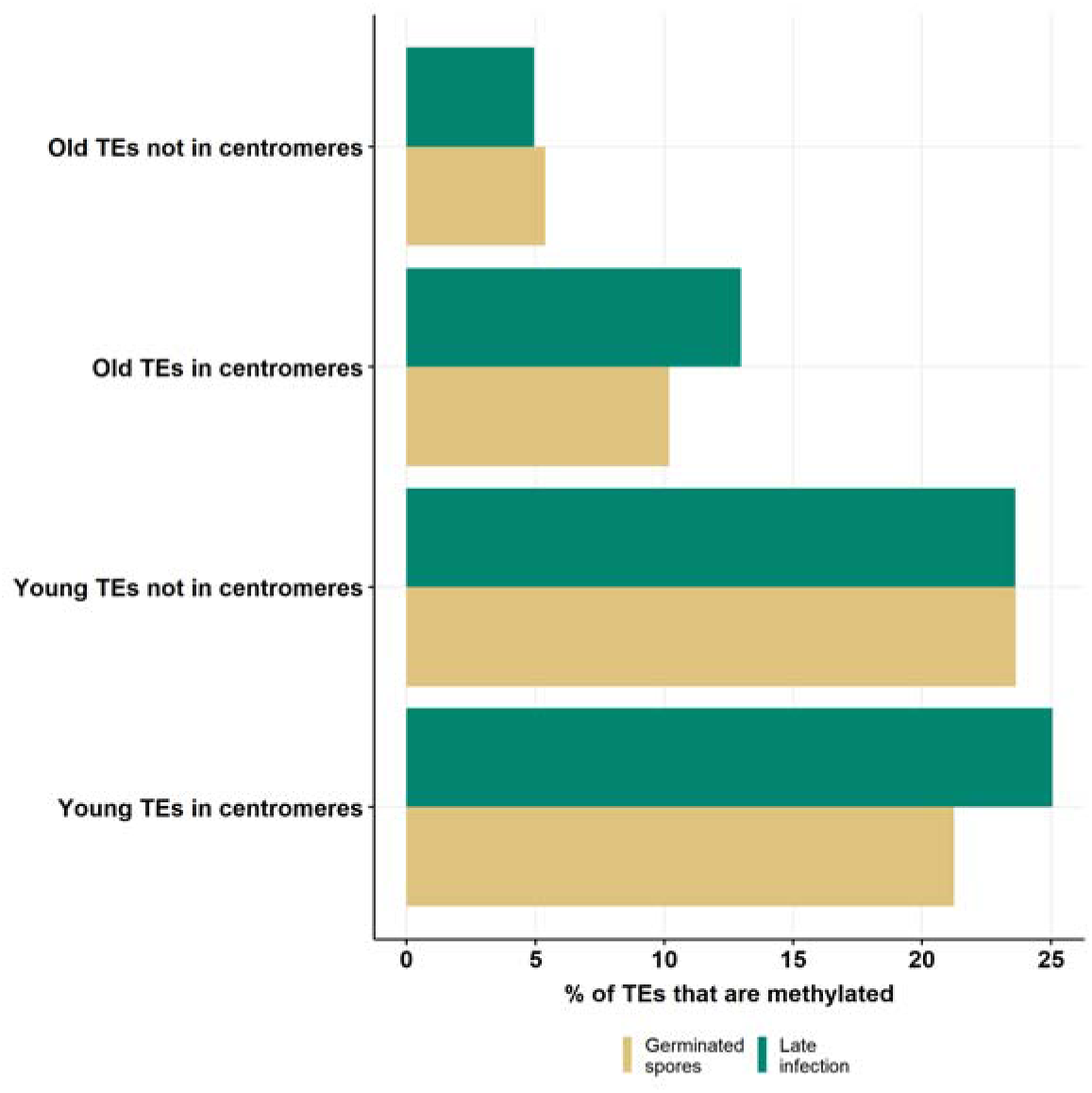
Proportions of young and old *Pgt* TEs that are methylated in the centromeres and outside the centromeres. Both inside and outside the centromeres, young TEs (> 90% family identity) are preferentially targeted by CG methylation.

### *Pgt* induces early and late waves of sRNAs with opposing profiles

To assess the role of the RNAi machinery in methylation, we performed sRNA-sequencing on germinated spores, uninfected wheat and infected wheat at 3 dpi, 5 dpi and 7 dpi. Adapter-trimmed and tRNA/rRNA-filtered sRNA reads were first mapped to the wheat and *Pgt* genomes. Strikingly, the read alignment rates show a strong presence of *Pgt* sRNAs in the late infection sample (7 dpi, Table 2). The mapping rates to rust in early infection (3 dpi and 5 dpi) are low at 5.25% and 5.37%, respectively, but increase drastically to 50.18% in late infection (7 dpi). In contrast, ∼67% of sRNA reads map to the wheat genome in early infection and in late infection the sRNA mapping rate to wheat decreases to 30.3%.

**Table 2:** Small RNA read mapping rates to the wheat and rust genomes.

| Sample | Number of reads | Mapped to *Pgt* | Mapped to wheat |
| --- | --- | --- | --- |
| Germinated spores | 27,536,477 | 55.93% | 0.73% |
| Uninfected wheat | 2,353,359 | 3.56% | 70.34% |
| Infected wheat 3dpi | 3,040,002 | 5.25% | 67.56% |
| Infected wheat 5dpi | 2,914,397 | 5.37% | 66.98% |
| Infected wheat 7dpi | 5,815,521 | 50.18% | 30.3% |

We predicted 7,304 *Pgt* sRNA loci (7,299 siRNAs and 5 miRNAs) and 411 wheat sRNA loci (360 siRNAs and 51 miRNAs) (Supplementary Files S1-S4). For each predicted sRNA locus, we obtained the most abundant sRNA. For predicted miRNA loci, this will generally be the functional mature miRNA. The read length distributions of rust and wheat sRNAs show different patterns and distinct peaks of abundance (Figure 7). The *Pgt*-derived sRNAs are predominantly 20, 21 or 22 nts in length. This is true for both for the single most abundant sRNA in each locus as well as for the total sRNA reads derived from each locus (Figure 7). There are two distinct peaks at 21 nt and 24 nt for the wheat sRNAs, as is expected for plant sRNAs. Most predicted wheat miRNAs are 21 nt and have a 5’ uracil (71.4%) while the wheat siRNAs are mostly either 21 nt with a 5’ uracil or 24 nt with a 5’ adenine (Table 3). The two distinct peaks at 21 and 24 nts with their corresponding 5’ nucleotide preferences support the predicted presence of both miRNAs and siRNAs in the wheat sRNA set and the 24 nt wheat siRNAs are likely involved in RdDM [8, 31]. However, two distinct classes of siRNAs also appear to be present in *Pgt* based on 5’ nucleotide preference, although differing in size to the wheat siRNAs. *Pgt* siRNAs of length 20-21 nts have a strong preference for a 5’ uracil (∼76%), whereas 54% of the 22 nt *Pgt* siRNAs have a 5’ adenine, suggesting functional diversification.

**Table 3:** Predicted miRNAs and siRNAs in *Pgt* and wheat and their properties.

|  | *Pgt* siRNAs | *Pgt* miRNAs | Wheat siRNAs | Wheat miRNAs |
| --- | --- | --- | --- | --- |
| # of sRNAs | 7,299 | 5 | 360 | 51 |
| 20 nts (5' A\| 5' U) | 26.5% (16% \| 76%) | 0% | 8.3% (17% \| 73%) | 13.7% (0% \| 100%) |
| 21 nts (5' A\| 5' U) | 34.7% (19% \| 77%) | 0% | 47.2% (21% \| 46%) | 68.6% (11% \| 71%) |
| 22 nts (5' A\| 5' U) | 31.8% (54% \| 44%) | 100% (60% \| 40%) | 14.7% (30% \| 60%) | 15.7% (0% \| 88%) |
| 24 nts (5' A\| 5' U) | 0% | 0% | 25.6% (45% \| 23%) | 0% |

**Figure 7:**
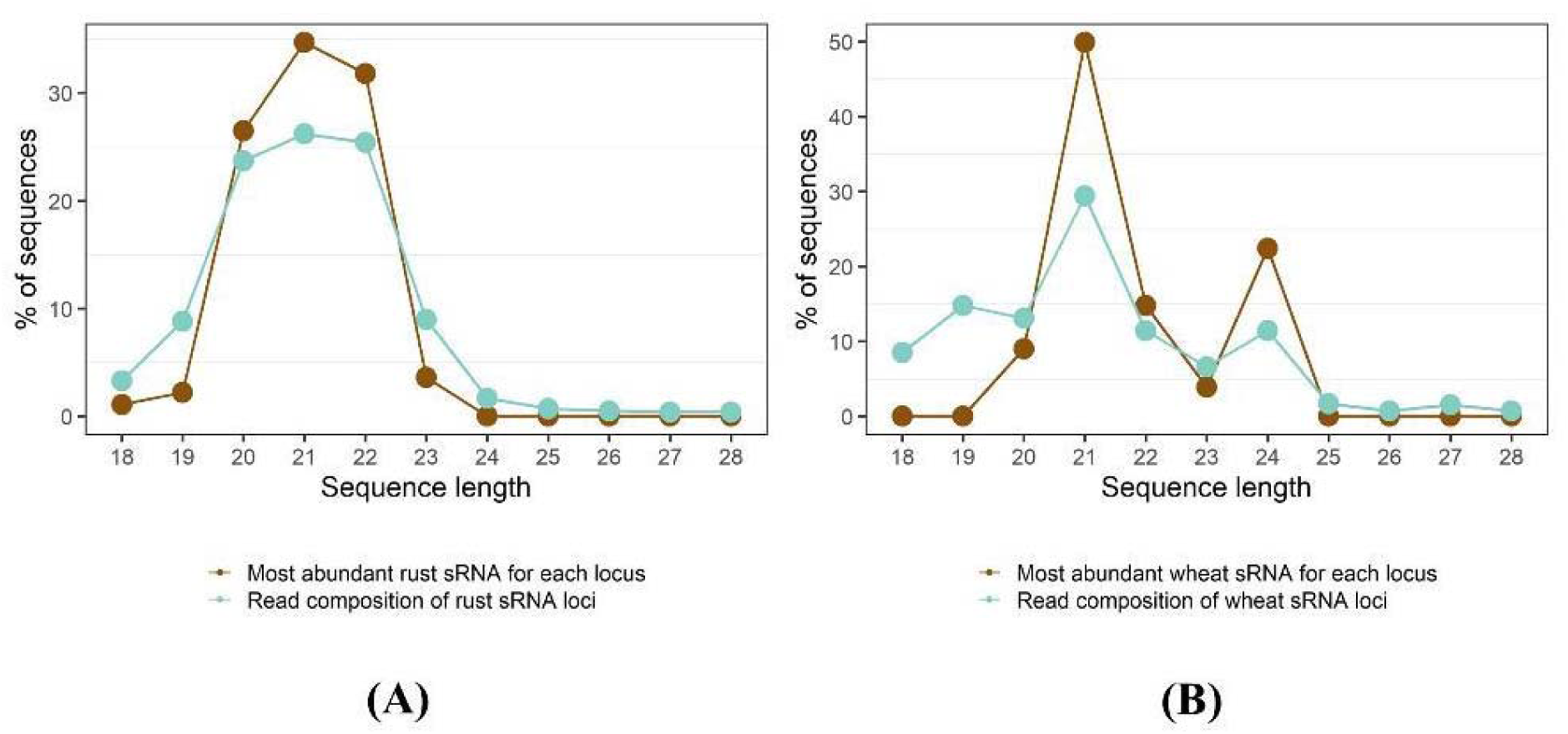
Sequence length distributions of predicted sRNAs in *Pgt* and wheat. **(A)** The rust sRNAs are predominantly 20-22 nt in length, whereas the **(B)** wheat sRNAs show strong peaks at 21 nt and 24 nt. Both the single most abundant RNA in each locus as well as the total reads forming the loci show the same peaks.

Next, we assessed the differential expression of *Pgt* sRNAs at the start of infection (germinated spores), during early infection (3 dpi and 5 dpi) and during late infection when sporulation begins (7 dpi). We detected no differential expression of *Pgt* sRNAs between 3 dpi and 5 dpi, likely due to the low number of mapped reads (Table 2, Figure 8A) and therefore combined these time points to represent early infection. Strikingly, 91.6% of the *Pgt* sRNA clusters are predicted as differentially expressed among germinated spores, early infection (3 and 5 dpi) and late infection (7 dpi): 2,714 are up-regulated in germinated spores, 509 up-regulated during early infection and 3,987 up-regulated during late infection (Figure 8, Supplementary Files S5-S8). A large proportion of the up-regulated sRNAs at the late infection time point (76.1%; 3,035 of 3,987) showed up-regulation against all the other conditions (germinated spores, 3 dpi and 5 dpi). In contrast, among the up-regulated sRNAs in germinated spores, the majority (87.5%; 2,377 of 2,714) are up-regulated against the late infection sample, with only a small number (33 and 35) being up-regulated compared to the 3 dpi or 5 dpi samples. Thus, the sRNAs up-regulated during late infection are highly specific to that time point, indicating the presence of early (germinated spores, 3 dpi and 5 dpi) and late (7 dpi) waves of *Pgt* sRNAs during wheat infection. In contrast to *Pgt*, which exhibits prominent early and late infection waves of sRNAs, only 14 of the 411 wheat sRNAs (3.4%) are predicted to be differentially expressed. Amongst these 14 differentially expressed wheat sRNAs there is no predicted miRNA.

**Figure 8:**
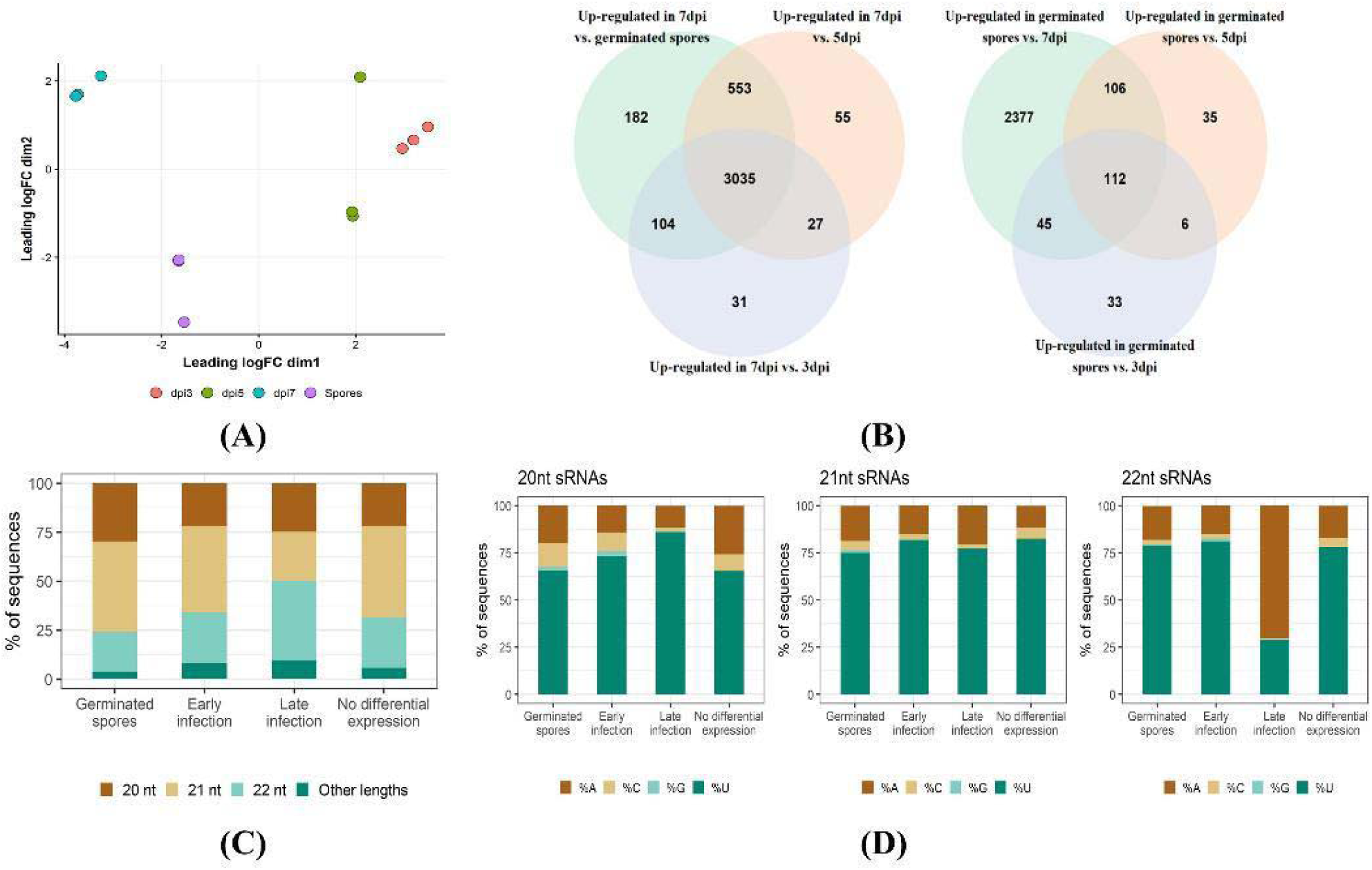
*Pgt* sRNA differentially expression analysis. **(A)** A multi-dimensional scaling plot using the edgeR package shows the clustering of the replicates for the different samples. The 3 dpi and 5 dpi samples show less differences in expression than the germinated spores and 7 dpi samples. **(B)** Venn diagrams of up-regulated *Pgt* sRNAs shared between the different time points: germinated spores, early infection (3 dpi and 5 dpi) and late infection (7 dpi). Two major classes of sRNAs occur: one that is up-regulated during late infection (*n* = 3,035) and one that is up-regulated in germinated spores compared to late infection (*n* = 2,377). **(C)** Sequence lengths and **(D)** 5’ nucleotide distribution of the *Pgt* sRNAs. *Pgt* sRNAs up-regulated during late infection differ in length distribution and 5’ nucleotide preference to the remaining *Pgt* sRNAs. 22 nt *Pgt* sRNAs up-regulated during late infection strongly prefer a 5’ adenine, which is not observed for 22 nt sRNAs expressed in the other conditions.

We assessed the length distributions and 5’ nucleotide preferences of differentially expressed *Pgt* sRNAs (Figure 8C, D). The early wave *Pgt* sRNAs are predominantly 21 nts in length (44% and 46.2%, respectively). In contrast, the largest class (40.8%) of the late wave *Pgt* sRNAs are 22 nts in length. *Pgt* sRNAs with no detected differential expression follow a similar size distribution pattern to the early wave sRNAs, with 21 nt sRNAs being the most prevalent class (46.4%, Figure 8C). The majority (60-80%) of the 20, 21 and 22 nt sRNAs up-regulated in germinated spores, during early infection and those with no differential expression contain a 5’ uracil (Figure 8D). This is also true for 20 and 21 nt late wave sRNAs. In contrast, the 22 nt late wave sRNAs have a strong preference for 5’ adenines (∼70%, Figure 8D). This suggests the specific induction of a different functional class of sRNAs during these late infection stages, similar to the occurrence of 24 nt siRNAs with a 5’ adenine in plants.

### Late wave *Pgt* sRNAs are produced from the centromeric regions, whereas the early wave sRNAs are highly conserved and derived from genes

The late wave *Pgt* sRNAs also exhibit opposing genomic origins to the early wave sRNAs (Table 4). The early wave sRNAs predominantly map to annotated genes (76.5% in germinated spores; 70.1% at 3 and 5 dpi), compared to only 16.5% of late wave sRNAs. Late wave sRNAs are largely generated from repetitive elements (86.9%), in contrast to the early wave sRNAs (24.2% in germinated spores and 28.7% at 3 and 5 dpi). Most of the repetitive elements associated with sRNAs belong to the class of LTR retrotransposons, particularly Gypsy elements. Strikingly, 24% of the late wave sRNAs originate from the centromeric regions, in contrast to only 1-3% of the early wave sRNAs and the sRNAs with no differential expression (Table 4).

**Table 4:** Genomic origins of *Pgt* sRNAs. The *Pgt* sRNAs map in similar proportions to the two haplotypes. More than half of sRNAs are conserved and have a homologous counterpart. Late wave sRNAs preferentially originate from repetitive regions and the centromeres.

|  | Early wave sRNAs |  | Late wave sRNAs | No differential expression |
| --- | --- | --- | --- | --- |
|  | Up-regulated in germinated spores | Up-regulated during early infection | Up-regulated during late infection |  |
| # of sRNAs | 2,714 | 509 | 3,987 | 617 |
| Centromeric sRNAs | 1.5% | 1.6% | 23.7% | 3.2% |
| On chromosomes A | 50.4% | 50.4% | 49.6% | 49.3% |
| On chromosomes B | 49.6% | 49.6% | 50.4% | 50.8% |
| sRNAs with homologous counterpart | 68.1% | 67.5% | 54.8% | 44.6% |
| Homologous counterpart is on alternate haplotype chromosome | 82.6% | 85% | 18.2% | 36.8% |
| Mapping to repeats | 25.1% | 30.3% | 87.9% | 62.4% |
| Mapping to genes | 76.5% | 70.1% | 16.5% | 47.2% |
| Overlap with methylated CG sites (late infection) | 5.5% | 10.6% | 53.8% | 27.9% |
| Overlap with methylated CG sites (germinated spores) | 6.0% | 12.6% | 45.1% | 25.8% |
| **Classification of repeats with mapped sRNAs** |  |  |  |  |
| **Class I (Retrotransposons)** | 59.1% | 56.6% | 56% | 60% |
| Gypsy LTR | 30.5% | 15.4% | 29.8% | 24.3% |
| Copia LTR | 9.9% | 18.3% | 14% | 13.5% |
| **Class II (DNA transposons)** | 37.8% | 38.9% | 42.7% | 39.5% |
| Tc1-Mariner | 5.8% | 5.7% | 2.9% | 4.4% |
| MuDR | 5.2% | 4.6% | 7.1% | 7.2% |

A gene function ontology (GO) term analysis of the 1,878 genes that are associated with *Pgt* sRNAs up-regulated in germinated spores reveals an enrichment in proteins with ATP binding activity as well as proteins with helicase and motor activity and RNA binding (Table 5). No significant enrichment in functional annotation was observed for genes that are associated with sRNAs with no differential expression, or with sRNAs up-regulated during early or late infection.

**Table 5:** *Pgt* genes that are associated with sRNAs up-regulated in germinated spores and their functional GO term enrichment. We assessed GO term enrichments of the annotated molecular function of *Pgt* genes that are associated with sRNAs compared to all other *Pgt* genes (FDR < 0.00001). The top ten categories with lowest FDR are shown.

| Enriched GO term category | False discovery rate (FDR) | # of genes |
| --- | --- | --- |
| **Genes that are associated with sRNAs up-regulated in germinated spores** |  |  |
| ATP binding | $1.7 \times 10^{-35}$ | 233 |
| Helicase activity | $1.5 \times 10^{-13}$ | 47 |
| Motor activity | $4.5 \times 10^{-11}$ | 25 |
| RNA binding | $6.5 \times 10^{-10}$ | 110 |
| Shikimate 3-dehydrogenase (NADP+) activity | $1.7 \times 10^{-9}$ | 8 |
| Histone methyltransferase activity (H3-K4 specific) | $4.2 \times 10^{-9}$ | 9 |

We further investigated the locations of the *Pgt* sRNAs on the 18 chromosome pairs and found that similar proportions occur in each of the two haplotypes (Table 4). We then assessed if sRNAs have a homologous counterpart on the corresponding haplotype. For this we re-mapped the sequencing reads that define a sRNA locus to the remainder of the genome and assigned the sRNA locus that has the highest coverage by those mapped reads as the homologous counterpart. Over two-thirds of sRNAs up-regulated in germinated spores have a homologous counterpart (68.1%, Table 4). Most of these homologous pairs (82.6%) are located on the corresponding chromosome from the alternative haplotype and generally occur in syntenous positions (Supplementary Figure S5). This is consistent with the observation that most of these sRNAs map to gene sequences which are expected to occur in allelic positions in each haplotype. In contrast, only around half of the late wave sRNAs have a homologous counterpart (54.8%), and only 18.2% of these homologous pairs are located on the corresponding chromosome (Table 4). In summary, the early wave sRNAs are conserved across the haplotypes and originate from gene models, whereas the late wave sRNAs originate from repetitive elements that are not conserved between haplotypes.

### During late infection, *Pgt* sRNAs are heavily transcribed from the centromeres and appear to direct genome-wide methylation to young TEs

To assess sRNA transcription in the centromeres, we plotted sRNA transcription levels at late infection and in germinated spores on the chromosomes. During late infection, strong peaks of sRNA transcription are apparent in the centromeric regions, except for chromosome 11A and 11B (Figure 9). The late wave *Pgt* sRNAs not only originate from the centromeric regions, they are also heavily induced from the centromeres in late infection. Whilst there is also transcription of sRNAs from the centromeres in germinated spores, the centromeric peak is less apparent and the overall transcription levels in the centromeres are only about a third of that observed during late infection (average reads per million: 160 at late infection and 54 in germinated spores).

**Figure 9:**
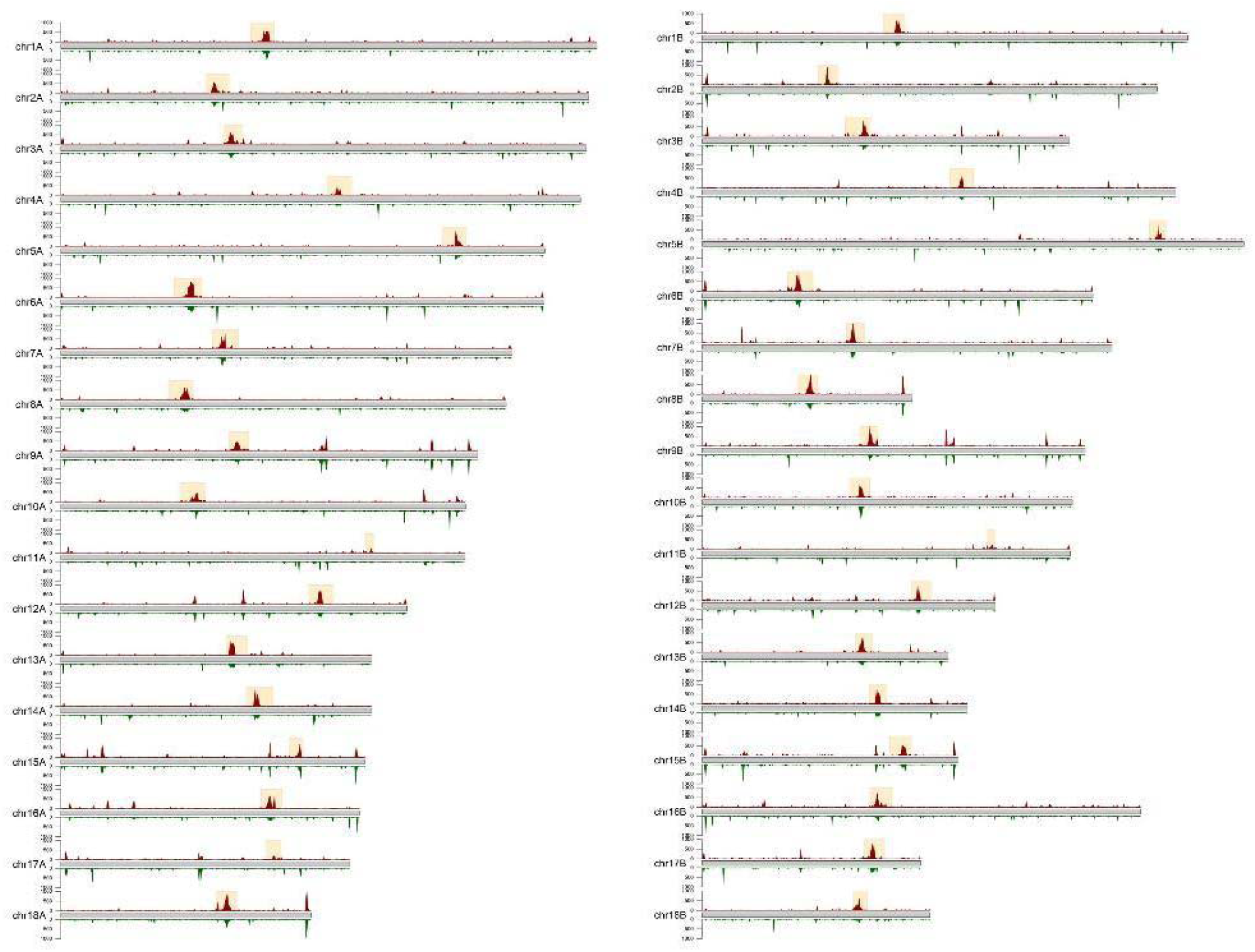
Transcription levels of sRNAs on the *Pgt* chromosomes. The transcription levels (reads per million per 10 kb genomic windows, < 1000 RPMs shown for clarity) of sRNAs are shown for late infection (red; above each chromosome) and for germinated spores (green; below each chromosome). Centromeric regions are indicated by yellow boxes. Higher transcription levels of sRNAs are seen from the centromeres during late infection than in germinated spores.

To investigate the genomic regions that might be targeted by these centromeric *Pgt* sRNAs, we re-mapped sRNAs without mismatches to the chromosomes and recorded all of their alignment positions. We then assessed which types of genomic regions are enriched for sRNA targeting. 30.8% of young TEs in the centromeres and 19.3% of the young TEs outside the centromeres have a late wave sRNA mapping to them, with much lower mapping rates to young TEs observed for the other sRNA classes (Table 6). This indicates that the late wave sRNAs might be involved in the silencing of young TEs, both inside and outside the centromeres.

**Table 6:** Young TEs preferentially overlap with late wave sRNAs both inside and outside the centromeres.

|  | Young TEs in centromeres | Old TEs in centromeres | Young TEs not in centromeres | Old TEs not in centromeres |
| --- | --- | --- | --- | --- |
| % that are sRNA+ (germinated spores) | 3.7% | 0.7% | 3.4% | 0.5% |
| % that are sRNA+ (early infection) | 2.1% | 0.3% | 1.7% | 0.2% |
| % that are sRNA+ (late infection) | **30.8%** | **7.0%** | **19.3%** | **1.8%** |
| % that are sRNA+ (no DE) | 7.1% | 0.5% | 5.2% | 0.4% |

**Table 7:** RNAi and 5mC methyltransferase genes in *Pgt*. For each protein, the identifiers of the allelic proteins on each haplotype are given. Homologs of the *Pgt* p7a PGTG_13081 and PGTG_13088 dicer proteins were not found in the gene annotation of *Pgt* 21-0.

| Gene annotation | Identifier | *Pgt* 21-0 proteins | *Pgt* p7a identifier |
| --- | --- | --- | --- |
| **Argonautes** | Argonaute 1 | PGT21_021399 (chr14A) and PGT21_022388 (chr14B) | PGTG_08429 |
|  | Argonaute 2 | PGT21_001976 (chr13A) and PGT21_002123 (chr13B) | PGTG_11327 |
| **Dicer** | Dicer 1 | PGT21_033256 (chr4A) and PGT21_033881 (chr4B) | PGTG_19535 |
|  | Dicer 2 | PGT21_033021 (chr4A) and PGT21_033709 (chr4B) | PGTG_19538 |
|  | Dicer 3 | PGT21_029367 (chr6A) and PGT21_028061 (chr6B) | PGTG_12289 |
|  | Dicer 4 | - | PGTG_13081 |
|  | Dicer 5 | - | PGTG_13088 |
| **RdRPs** | RdRP 1 | PGT21_002642 (chr10A) and PGT21_001684 (chr10B) | PGTG_20838 |
|  | RdRP 2 | PGT21_009430 (chr15A) and PGT21_009102 (chr15B) | PGTG_17766 |
|  | RdRP 3 | PGT21_009651 (chr14A) and PGT21_011158 (chr14B) | PGTG_02834 |
|  | RdRP 4 | PGT21_031631 (chr4A) and PGT21_032301 (chr4B) | PGTG_05092 |
|  | RdRP 5 | PGT21_031875 (chr8A) and PGT21_035256 (chr16B) | PGTG_09533 |
| **5mC MTases** | DNMT1 | PGT21_014413 (chr4B) and PGT21_012711 (chr4A) | PGTG_03742 |
|  | DNMT5 | PGT21_036642 (chr1A) and PGT21_037052 (chr1B) | PGTG_17071 |

To address this further, we explored whether a TE that has a sRNA mapping to it (TE^sRNA+^) is more likely to be methylated than a TE that does not have a sRNA mapping to it (TE^sRNA-^). Indeed, we observed that the late wave *Pgt* sRNAs are strongly associated with methylation of TEs, particularly young TEs, both inside and outside the centromeres (Figure 10). In late infection, 33.2% of TEs^sRNA+^ are methylated if the overlapping sRNAs are late wave sRNAs. In contrast, only 8.3% TEs^sRNA+^ are methylated if the overlapping sRNAs have no differential expression. Whilst the late wave sRNAs are preferentially 22 nts in length with a 5’ adenine (Figure 8), only 16.7% of TEs^sRNA+^ are methylated if they overlap with this subclass class of late wave sRNAs. This suggests that the up-regulation at late infection drives RNA-directed methylation in *Pgt*, and is not only restricted to the 22 nt sRNAs and a 5’ adenine.

**Figure 10:**
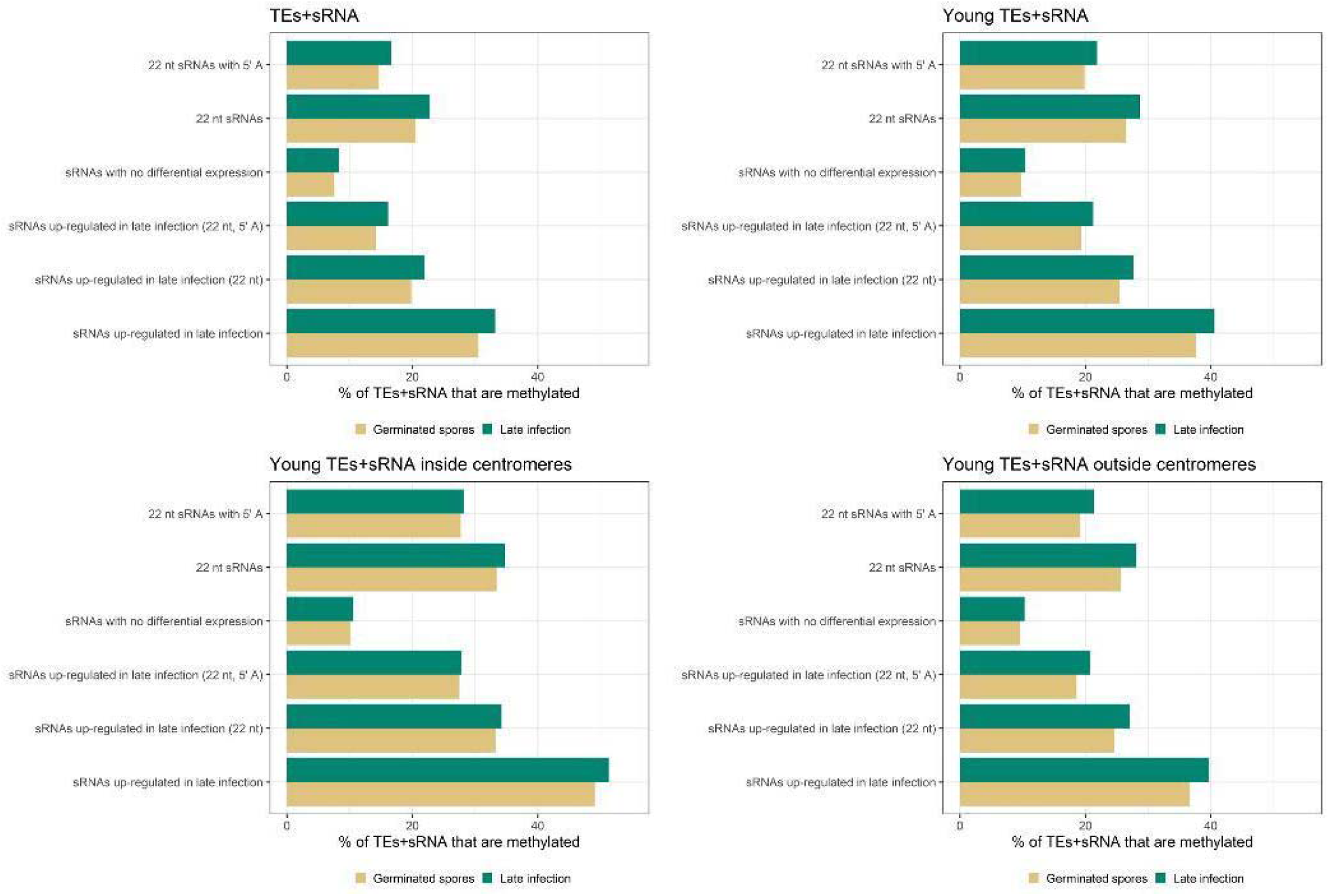
Percent of TEs that are methylated in germinated spores and late infection and overlap with sRNAs (TEs^sRNA+^). TEs both inside and outside the centromeres that overlap with late wave sRNAs (TEs^sRNA+^) are more likely to be methylated, suggesting that *Pgt* is using sRNA-directed methylation.

Young TEs both inside and outside the centromeres appear to be particularly targeted for RNA-directed methylation by late wave sRNAs (Figure 10). Inside the centromeres, 51.3% of the TEs^sRNA+^ are methylated in late infection when they overlap with late wave sRNAs. Strikingly, RNA-directed methylation is also pronounced outside the centromeres where 39.7% of the TEs^sRNA+^ are methylated in late infection when they overlap with late wave infection sRNAs. Taken together, our results indicate that the late wave *Pgt* sRNAs originate mainly from the centromeres but seem to direct DNA methylation to loci homologous to their sequences both inside and outside the centromeres, preferentially targeting young TEs.

### TE-associated CG methylation leads to silencing of nearby genes

We investigated the effect of CG methylation and sRNA-directed CG methylation on overlapping or adjacent genes. 4,650 *Pgt* genes overlap with CG methylation sites in germinated spores and 4,848 genes overlap with CG methylation sites in late infection. The proteins encoded by these methylated genes are not enriched for secreted proteins or for any GO terms (data not shown). Almost all methylated genes overlap with repeats (except for 363 genes in germinated spores and 428 in late infection), suggesting that CG methylation predominantly correlates with TE silencing in *Pgt*.

To determine if the presence of methylated TEs affects nearby gene expression, we assessed transcription levels of genes that are close to or overlap with TEs. For both germinated spores and infected leaves, genes that have an overlapping methylated TE have significantly lower gene transcription levels than those overlapping with a non-methylated TE (Figure 11). This silencing effect is also observed for genes that are close (< 500 bps) to a methylated TE compared to genes that are close to a non-methylated TE (Figure 11A), while no difference was observed for genes > 500 bps from methylated TEs. We then compared the expression levels of genes that overlap with TEs following whether they also overlap with the late wave sRNAs or not. Whilst genes that overlap with methylated TEs have low expression levels in general, they have significantly lower levels of expression when the TE is additionally targeted by a late wave sRNA and this holds true for both young and old TEs (Figure 11B).

**Figure 11:**
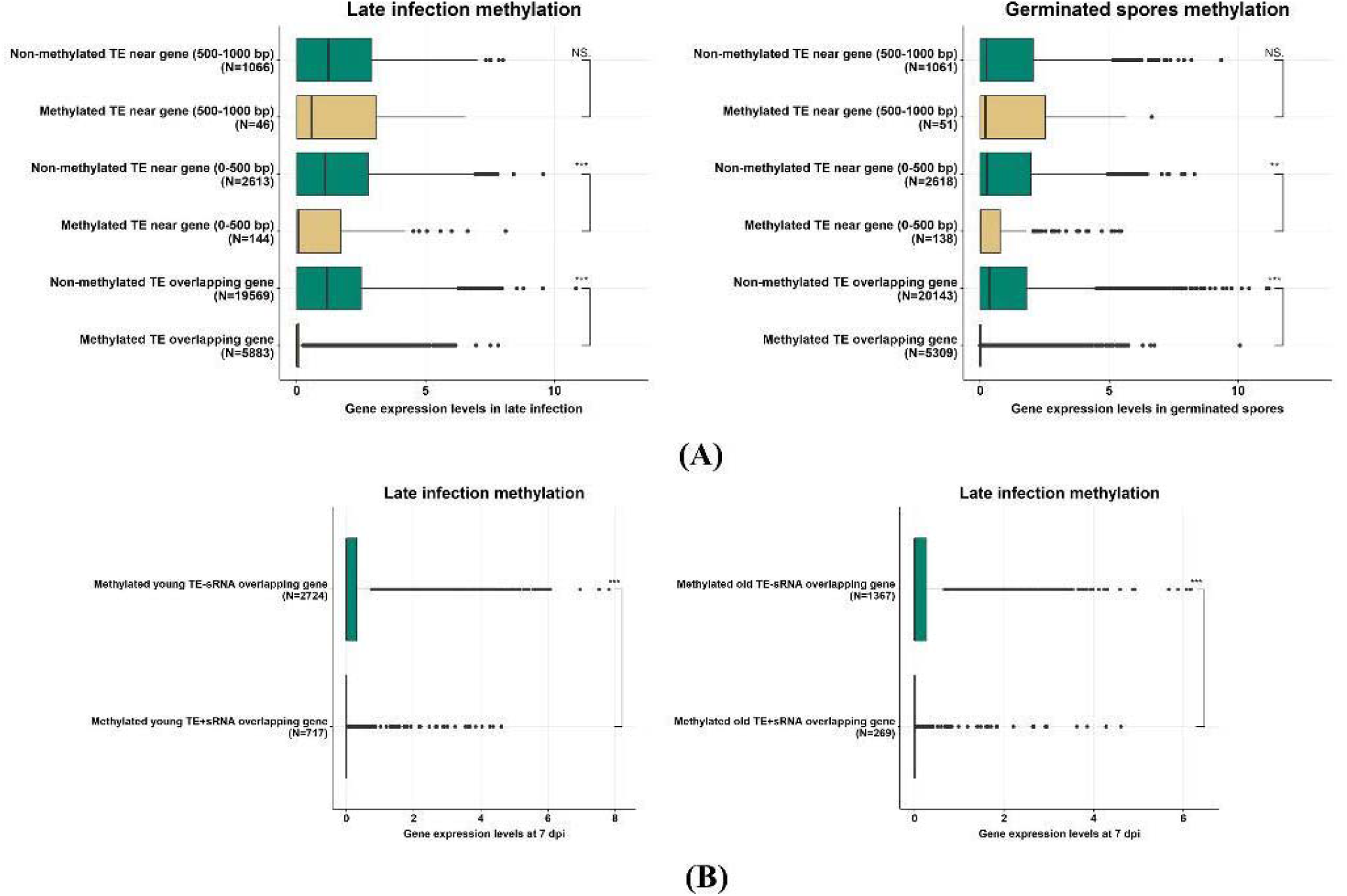
Expression levels of genes that have an inserted methylated repeat or that overlap with methylated repeats. **(A)** Expression levels (*log*-normalized transcripts per million) of genes are shown. A strong silencing effect is shown for genes that contain a methylated repeat, both in germinated spores and during late infection. Genes that are near a methylated repeat (up to 500 bps) also show suppressed expression. **(B)** This gene silencing is more pronounced if a TE has a late wave sRNA mapping to it (TEs^sRNA+^), both for young and old TEs.

### A subset of RNAi genes are up-regulated during late infection, supporting functional diversification

RNAi machinery genes were previously identified in the reference genome *Pgt* p7a [27, 32]. We searched for the *Pgt* p7a RNAi genes in the gene annotation of the fully phased, chromosome-scale assembly of *Pgt* 21-0. Two argonaute genes, three dicer genes and five RdRP genes are present in the annotation of *Pgt* 21-0 on each haplotype (Table 8). We then searched for 5mC methyltransferase (5mC MTase) genes. Four classes of fungal DNA methyltransferases have been observed in fungi, but basidiomycetes predominantly have the DNMT1□and DNMT5 genes [33]. We identified DNMT1 and DNMT5 in the *Pgt* 21-0 annotation by searching for the previously identified *Pgt* p7a genes [4] and additionally for the DNA methylase domain PFAM domain (PF00145). In line with Bewick AJ, Hofmeister BT, Powers RA, Mondo SJ, Grigoriev IV, James TY, Stajich JE and Schmitz RJ [4] we found homologs of the DNMT1□and DNMT5 genes and confirmed the absence of the other two classes (DIM-2 and RID) and of 6mA DNA and RNA MTase genes in *Pgt* 21-0. The lack of 6mA DNA and RNA MTase genes in *Pgt* indicates that cytosine methylation is the primary DNA methylation process active in this species.

The gene expression profiles of the RNAi and 5mC MTase genes during a time course of *Pgt* 21-0 infecting wheat from 2-7 days post infection (dpi) [26] and in germinated spores and haustorial tissue [25] indicate two main patterns (Figure 12A): one set of RNAi genes (RdRPs 2/4/5, AGO2 and dicers 1/2) and 5mC Mtase genes that are constitutively expressed during infection, with the AGO2 genes showing particularly high expression; and another set of RNAi genes (AGO 1, dicer 3 and RdRPs 1/3) that are highly expressed only during the later stages of infection, with no or very low expression in germinated spores and during early infection. We did not observe differences in expression patterns of the RNAi genes between the two *Pgt* haplotypes.

**Figure 12:**
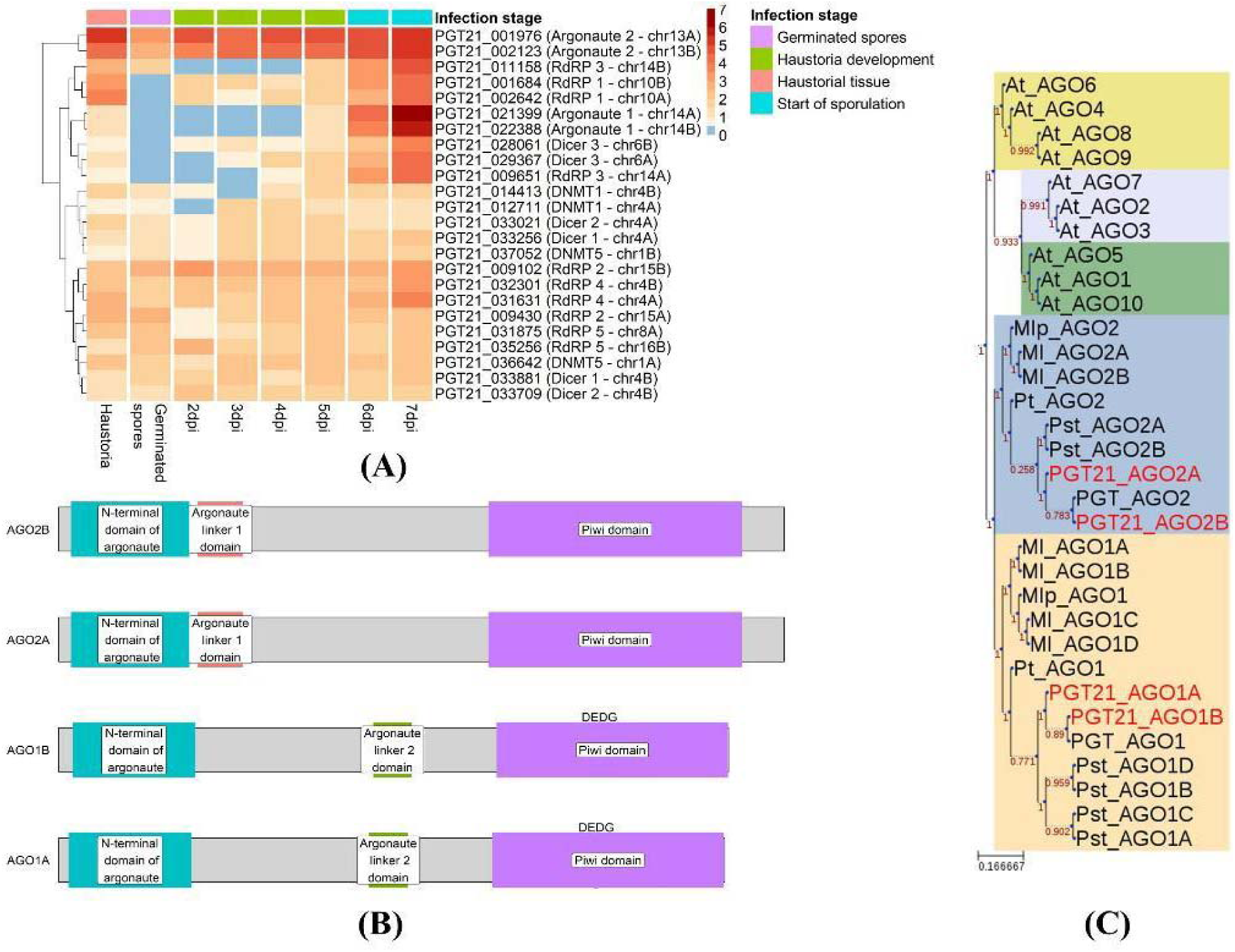
*Pgt* 21-0 RNAi and 5mC methyltransferase gene expression. **(A)** Hierarchical clustering of expression levels of *Pgt* RNAi genes in transcripts per million (*logTPM*, red color intensity relates to high expression). The *Pgt* RNAi RdRPs 1/3, argonaute 1 and dicer 3 show distinct high expression at sporulation in the later stages of infection (6-7 dpi). The 5mC methyltransferases DNMT1 and DNMT5 are expressed across all conditions. **(B)** The *Pgt* argonaute proteins have diversified on the sequence level and AGO 1 and AGO 2 show differences in protein domains. **(C)** A phylogenetic tree of *Arabidopsis* argonaute proteins (At_AGO1-10) and other rust argonautes (Mlp: *Melampsora larici-populina*; Ml: *Melampsora lini*; Pt: *Puccinia triticina*; Pst: *Puccinia striiformis* f. sp. *tritici;* PGT: *Puccinia graminis* p7a) supports the diversification of cereal rust argonautes into two classes, AGO1 and AGO2.

A protein domain analysis further supports the functional diversification of the *Pgt* argonautes AGO 1 and 2. AGO 1 has an argonaute linker 1 domain and is longer in sequence, whereas AGO 2 has an argonaute linker 2 domain (Figure 12B). A phylogenetic tree constructed from the argonaute proteins of *Arabidopsis thaliana* and several rust fungi further supports the diversification of the rust AGOs into two classes. In *Arabidopsis thaliana*, AGO1 and AGO10 bind preferentially small RNAs with a 5′ uracil, whereas AGO2, AGO4, AGO6, AGO7 and AGO9 prefer sRNAs with 5’ adenines and AGO5 5’ cytosines [1] and their diversification into these three classes is apparent from a phylogenetic tree (Figure 12C). The rust argonautes are distinct from the *Arabidopsis* clade and divide into two distinct families, with one copy of each present in the haploid genomes of each rust species. Taken together, the expression and sequence analyses show that *Pgt* RNAi machinery has functionally diversified and suggests that *Pgt* might use RNAi to regulate stage-specific infection processes, such as during the formation of new urediniospores during late infection.

## Discussion

Epigenetic silencing mechanisms mediated by sRNAs and methylation are not well-studied in plant-pathogenic fungi [10] and had thus far not been described in rust fungi. Current knowledge has been derived mainly from model species which comprise a relatively small group of fungi. Through sRNA sequencing over a time course of *Pgt*-wheat infection, we uncovered that *Pgt* produces two distinct waves of sRNAs with different profiles during infection and over 90% of its sRNAs are differentially expressed. Previous studies on sRNA characterization in fungal plant pathogens mostly rely on sequencing of one time point of infection, which obscures the expression profiles of sRNAs over time. For example, a study in the stripe rust fungus *Puccinia striiformis* f.sp. *tritici* sequenced sRNAs at 4 dpi and found that the majority of the predicted 20–22 nt *Pst* sRNAs carry a 5’ uracil [34]. The presence of distinct sRNA profiles in mycelia and appressoria tissues was suggested in the rice blast fungal pathogen, *Magnaporthe oryzae* [16]. However, prominent waves of sRNA expression profiles during infection of plants had thus far not been reported.

*Pgt* sRNA expression is under tight temporal control, with ∼90% of *Pgt* sRNAs differentially expressed over the time course. The presence of two distinct sRNA profiles has thus far not been observed in rust fungi and supports functional diversification of the RNAi machinery, with a strong role in the infection and proliferation process. The early wave sRNAs are predominantly 21 nts with a 5’ uracil derived from genes. In contrast, the late wave sRNAs are mainly 22 nt sRNAs with a 5’ adenine derived from repetitive sequences. We speculate that the majority of 22 nt *Pgt* sRNAs are responsible for transcriptional silencing of TEs during sporulation and the majority of 20-21 nt *Pgt* sRNAs for posttranscriptional silencing of genes. This is similar to what has been reported in plants, which produces 20-22 nt miRNAs/siRNAs and 24 nt heterochromatic sRNAs [35]. In plants, TEs are silenced mainly via 24 nt sRNAs in the RdDM pathway [1]. These 24 nt sRNAs are most abundant during seed development in plants, presumably to ensure stable inheritance of the genome.

The up-regulation of 22 nt *Pgt* sRNAs with enrichment for 5’ adenines during late infection coincides with the up-regulation of the AGO1 gene. Similarly, the preferential accumulation of 21 nt 5’ uracil sRNAs in germinated spores and during early infection correlates with high-level expression of AGO2 and relatively low expression of AGO1. This suggests that similarly to plants, the 5’ nucleotide of *Pgt* sRNAs might have a strong effect on preferential loading into different argonautes. In *Arabidopsis thaliana*, AGO1 and AGO10 bind preferentially small RNAs with a 5′ uracil, whereas AGO2, AGO4, AGO6, AGO7 and AGO9 prefer sRNAs with 5’ adenines and AGO5 5’ cytosines [1]. Our analysis suggests that *Pgt* AGO2 preferentially loads sRNAs with a 5’ uracil and AGO1 preferentially binds 22 nt sRNAs with a 5’ adenine, which is worthy of investigation in future experimental studies.

We discovered striking parallels between *Pgt* sRNAs and plant sRNAs, in particular compelling evidence for a sRNA-directed TE silencing pathway in *Pgt* that resembles the RdDM pathway in plants. Such a RdDM-like pathway has thus far not been reported in fungi and might suggest that *Pgt* uses similar strategies to plants to maintain its highly repetitive genome [36]. The overlap of the late wave *Pgt* sRNAs with cytosine methylation sites suggests that these sRNAs may function similarly to plant 24 nt siRNAs to direct methylation to cause transcriptional silencing. The specific expression of one argonaute, one dicer and two RdRPs at the late stage of infection underlines their involvement in such a functionally diversified TE silencing pathway.

Furthermore, we showed that Hi-C data can be used to define centromeric regions in fungi and uncover the first centromeres in rust fungi. The *Pgt* centromeres are highly repetitive, hyper-methylated regions with exceptional sequence divergence, unexpectedly even between some haplotypes. Highly repetitive loci such as centromeres can generate sRNAs which in turn are required for epigenetic silencing [37]. Centromeres are essential for chromosome segregation during cell division and heterochromatin is vital to maintain the integrity of the centromeres. Eukaryotic centromere sequences are highly diverse in sequence and can differ even between closely related species [38]. In fungi, their lengths range from point centromeres (<400 bp), short regional centromeres (>400 bp, <20 kb) to large regional centromeres (>20 kb) [39]. For example, the fission yeast *S. pombe* centromeres span between 35-110 kb and resemble those of vertebrates (central core domain of non-repetitive AT-rich DNA flanked by outer repeats), where the kinetochore is embedded in the heterochromatin of the outer repeats. In *Neurospora crassa*, centromeres are repetitive, AT-rich 150 to 300□kb long regions [40]. The human fungal pathogen *Cryptococcus* harbours large regional centromeres that are rich in LTR retrotransposons [18]. The formation of silent heterochromatin in some yeasts depends on siRNAs derived from pericentromeric regions and on the RNAi machinery [12, 41]. Genes placed near centromeric chromatin are typically silenced [42, 43], with the strongest repression at the outer repeats [44, 45]. In the rice blast fungus *Magnaporthe oryzae*, centromeres span 57-kb to 109-kb transcriptionally poor regions and share highly AT-rich and heavily methylated DNA sequences [46]. Clearly, centromeres are not well-studied in plant-pathogenic fungi and had thus far not been described in rust fungi. The high activity of *Pgt* centromeric sRNAs in the later stages of infection might ensure that the genome is passed on stably to subsequent generations through methylation and condensation of centromeres. The TE silencing function can have a silencing effect on nearby genes, and this seems to occur in some *Pgt* genes that overlap with methylated TEs. In plants, insertion of TEs near genes can provide cis-elements for stress responsive or tissue-specific expression, and the expression level can be modulated by DNA methylation and/or histone modification at the TEs due to siRNA targeting. It is likely that a similar DNA methylation or histone modification mechanism exists in *Pgt*.

In contrast to plants, the roles of sRNAs in epigenetic silencing pathways of fungal plant pathogens has been understudied and previous research has focused heavily on the roles of sRNAs in cross-kingdom gene silencing [47, 48]. Several cross-kingdom RNAi interactions between fungal pathogens and plants have been uncovered. Some *Botrytis cinerea* sRNAs silence *Arabidopsis* and tomato genes involved in plant immunity and are mainly derived from LTR retrotransposons and are 21 nt in size with a 5’ uracil [49], while *Arabidopsis* cells secrete exosome-like extracellular vesicles to deliver sRNAs into the fungal pathogen *Botrytis cinerea* to silence pathogenicity genes [50]. A wheat stripe rust fungus *Puccinia striiformis* f. sp. *tritici* 20 nt sRNA has been suggested to target the wheat defence pathogenesis-related 2 (*PR2*) gene [51]. The fungal pathogen *Sclerotinia sclerotiorum* produces mainly 22-23 nt sRNAs with a 5’ uracil from repeat-rich regions during infection [52]. Whilst *Pgt* might also use sRNAs to target host genes for silencing, we found strong support for endogenous roles of *Pgt* sRNAs during infection. Using the ShortStack software which uses criteria tailored to plant miRNA properties, we predicted only a handful of *Pgt* sRNAs that fulfil the criteria for miRNAs and thus might represent sRNAs involved in gene silencing. However, it is possible that *Pgt* produces a larger contingent of miRNA-like RNAs that follow currently unknown fungal-specific rules. Loci with some, but insufficient, evidence for miRNA biogenesis (such as strandedness) using the ShortStack software might be worth exploring as miRNA-like candidates in the future [53]. We did not perform target prediction of *Pgt* sRNAs due to the lack of fungal-specific targeting rules and the high false positive rate of miRNA target prediction tools [54]. In future studies, sRNA-sequencing specifically of haustorial tissues can help to elucidate if haustoria are potentially sites of sRNA transfer between the host and rust fungi [55] and then we can combine target prediction with gene expression data to reduce the number of false positive predictions.

## Conclusions

The wheat stem rust disease caused by *Puccinia graminis* f. sp. *tritici* (*Pgt)* is one of the most devastating crop diseases and of significant global interest. Our work uncovers fundamental characteristics of the stem rust RNAi machinery, DNA methylation in rust fungi and the first characterization of centromeres in rust fungi. We found compelling evidence for an sRNA-directed DNA methylation pathway in rust fungi, similarly to the RdDM pathway in plants. *Pgt* induces waves of early and late infection sRNAs with differing profiles and up-regulates a subclass of RNAi genes during late infection. Future research can use this knowledge to optimize methods of host-induced gene silencing where sRNAs from the plant operate via the fungus’s own RNAi machinery to silence pathogen genes important for causing disease.

## Methods

### Hi-C data analysis and centromere identification

Previously published Hi-C data [24] available in NCBI under BioProject PRJNA516922 was analyzed using HiC-Pro 2.11.1 [56] and contact maps were plotted with HiCExplorer’s hicPlotMatrix [57] to identify centromeric regions.

Chromosomes and centromeric regions were aligned using DGenies [58] and regions of macrosynteny were extracted from the minimap2 [59] paf alignment produced by DGenies. Pairwise *k*-mer distance estimations were calculated using Mash 2.2.0 with the function mash triangle [60] and clustered as a dendogram (hclust with the method ward.D2).

### Gene expression analysis and repetitive element annotation

Previously published RNA-seq data (0 dpi, 2 dpi, 3 dpi, 4 dpi, 5 dpi, 6 dpi, 7dpi) was used for the gene expression analysis [26]. This was complemented with previously published RNA-sequencing data of *Pgt* 21-0 germinated spores and haustorial tissue [61]. We used Salmon 1.1.0 to align reads to the *Pgt* 21-0 transcripts [24] and to estimate transcript abundances in each sample (salmon index –keepDuplicates and salmon quant –validateMappings). We used tximport and DESeq2 to assess gene differential expression [62, 63]. Differentially expressed genes were annotated with the B2GO software and GO term enrichment analyses were performed with B2GO and the category molecular function [64].

Transcription levels on the chromsomes were obtained by aligning the RNA-seq reads to the *Pgt* chromosomes [24] with HISAT2 2.1.0 and default parameters [65]. Bedtools 2.28.0 was used to slice the chromosomes into windows (bedtools makewindows) and the aligned reads per genomic window were counted (bedtools coverage – counts) and normalized to reads per million.

Repeat regions were annotated as described previously [66, 67] using the REPET pipeline v2.5 [29, 68, 69] for repeat annotation in combination with Repbase v21.05 [70]. For *de novo* identification, we predicted repeats for both haplotypes independently using TEdenovo. We combined the resulting *de novo* repeat libraries without removing redundancies. We annotated both haplotypes with the combined TEdenovo repeat library and two additional repeat libraries from Rebase (repbase2005_aaSeq and repbase2005_ntSeq). We generated superfamily classifications as described previously [66, 67].

### Methylation sequencing and analysis

*Triticum aestivum* cultivar Rangcoo seeds were sown and stratified at 4°C with no light for approximately 48 h. To germinate, the pots were transferred to a growth cabinet set at 21°C with 60-70% relative humidity and a 16 h light cycle. Six days after sowing (seedling approximately 6 cm tall), plant growth inhibitor maleic hydrazide was added, 20 mL per pot at a concentration of 1.1 g/L. Infection of wheat seedlings was performed seven days after sowing. 300-400 mg of dormant *Pgt* urediniospores were heat-shock activated for approximately 3 min at 42-45°C. The urediniospores were then suspended in Novec™ 7100 solvent (3M™) and immediately used to inoculate the wheat seedlings, applying the suspension across leaves homogeneously using a flat paintbrush. Pots were then placed into a plastic container, leaves sprayed with Milli-Q water (Merck), sealed with a lid and placed in a secure transparent plastic bag. The bag was transferred to a growth cabinet set at 23°C with 60-70% relative humidity and a 16 h light cycle. After 48 h, the plants were removed from the bag. Infected leaves (day 7) were grinded in mortar and pestle using liquid nitrogen, converted into fine powder and lysed with the lysis buffer followed by DNA binding, ethanol wash and elution. Notable changes include the addition of 6 mM EGTA to the lysis buffer. DNA was further purified which included RNA/proteins removal, clean-up with cholorform:isoamyl alcohol (24:1), shearing with 5 passes through a 29 gauge needle and size selected with a Short Read Eliminator (SRE) XS 10 kb (Circulomics) according to dx.doi.org/10.17504/protocols.io.betdjei6.

To prepare material from germinated *Pgt* spores, freshly harvested spores (450-500 mg) were sprinkled on top of autoclaved Milli-Q (MQ) water in a glass baking tray and were incubated at 100% humidity at 20°C in dark for 18 h before harvesting. Once germination was assessed and verified by bright field microscopy, the layer of germinated spores was collected using a glass slide and the remaining moisture was removed as much as possible with paper towel. Dried sample were snap frozen in liquid nitrogen and stored at −80°C until processed for DNA extraction. High-molecular weight DNA was extracted from germinated spores following the Phenol:Chloroform method with minor modifications. Briefly, the germinated spores were grounded with liquid nitrogen into fine powdered material, suspended in lysis buffer followed by cholorform:isoamyl alcohol (24:1) steps twice, incubated in 200 µl of 10 mM Tris pH 8 and 200 µl of Tris-EDTA buffer (TE) at room temperature overnight. For secondary cleanup, DNA bound to Sera-Mag™ SpeedBead magnetic carboxylate-modified particles (GE Healthcare), washed 3 times with ethanol and eluted with 10 mM Tris-HCl pH 8. The DNA was size selected with a Short Read Eliminator (SRE) XS 10 kb (Circulomics) according to dx.doi.org/10.17504/protocols.io.betdjei6.

To perform native DNA sequencing, Oxford Nanopore Technologies (ONT) portable MinION Mk1B was adopted. Native DNA sequencing libraries were constructed according to the manufacturer’s protocol 1D genomic DNA by ligation (SQK-LSK109), using 3 µg of DNA input. Briefly, DNA was repaired (FFPE DNA Repair Mix, New England BioLabs® (NEB)), end-prepped with an adenosine overhang (Ultra II end repair / dA-tailing module, NEB), purified (AMPure XP, Beckman Coulter) and an ONT adapter was ligated each end (Quick T4 Ligation Module, NEB). Following, the library was cleaned once more and quantified using a Qubit Fluorometer (Thermo Fisher Scientific). A MinION FLO-MIN106 9.4.1 revD flow cell was primed, approximately 300 ng of library was loaded and sequenced according to the manufacturer’s instructions (ONT). When the majority of pores became inactive (approximately 24 h), the flow cell was treated with DNase I and another 300 ng of library was loaded, according to the ONT nuclease flush protocol. The nuclease flush protocol was performed 2-3 times, until the flow cell was expended.

### Bioinformatic processing and methylation calling

Raw fast5 reads were basecalled with Guppy version 3.4.5 (ONT), using the --fast5_out option. Sequencing output and quality was inspected with the NanoPack tool NanoPlot version 1.28.2 [71]. To identify *Pgt* reads, we mapped all Nanopore reads from both analyzed conditions against the *Pgt* genome [24] using minimap2 version 2.17-r941[59] evoking the nanopore flag (-map-ont). Only reads aligning to the *Pgt* 21-0 reference genome were used for downstream. *De novo* identification of the DNA modifications 5mC and 6mA was performed using Tombo version 1.5.1 [72]. We called 6mA and 5mC methylation with Tombo 1.5.1 following the github instructions. We converted resulting Bigwig files into Bed6 files by calling sites methylated that had per read methylation frequency above 0.5. We called CG methylation using nanopolish 0.12.3 and pycometh v0.4.2 (https://doi.org/10.5281/zenodo.3629254). A genomic element was considered methylated if at least two methylation sites mapped to it. Di- and trinucleotide frequencies were calculated with compseq from EMBOSS 6.6.0 [73].

### Small RNA sequencing, read processing, filtering and alignment

From the same infected leaf samples as the RNA-seq data [26], small RNA sequencing data was obtained. For rust infection, host plants (cv. Sonora) were grown at high density (∼25 seeds per 12cm pot with compost as growth media) to the two leaf stage (∼7 days) in a growth cabinet set at 18-23°C temperature and 16 h light. Spores (−80°C stock) were first thawed and heated to 42°C for 3 minutes, mixed with talcum powder and dusted over the plants. Pots were placed in a moist chamber for 24 hours and then transferred back to the growth cabinet. Leaf samples were harvested at specified days after inoculation, snap frozen and stored at −80°C until use. 100 mg freshly collected spores were germinated overnight in four 15 cm petri dishes, each containing 200ml sterile RO water. Germinated spores were harvested via filtering through nylon mesh 15 μm. Small RNAs were extracted from the germinated spores and infected leaf samples with the Purelink microRNA Isolation Kit from Invitrogen. We sequenced sRNAs (50 bp reads) from the following five conditions (3 replicates each) on the Illumina HiSeq: germinated spores, uninfected wheat and infected wheat at 3 dpi, 5 dpi and 7 dpi. Adapters were trimmed using cutadapt (-m18 –M28 -q30 –trim-n –discard-untrimmed) [74]. Untrimmed reads, reads shorter than 18 nts or reads larger than 28 nts were discarded and flanking N bases were removed from each read [74]. FASTQC was run on the resulting reads (http://www.bioinformatics.babraham.ac.uk/projects/fastqc/).

To eliminate reads derived from non-small RNAs, we first generated a database set of potential contaminating RNA sources. *Triticum aestivum* and *Puccinia* tRNAs, rRNAs and spliceosomal RNAs were collected from the RNACentral database [75] as well as the tRNA and rRNA RFAM families RF00001, RF00002, RF00005, RF01852, RF01960 and RF02543 [76], snoRNAs from dbsnOPY, 5S and 23S ribosomal RNAs from the European Nucleotide Archive (ENA) and the tRNA/rRNA file from the sRNA workbench [77]. This set of potential contaminant sequences was de-duplicated using bbmap and its tool dedupe.sh (sourceforge.net/projects/bbmap/). Reads that mapped to this set were removed using bowtie 1.1.2 [78]. To assess read length distributions across the different samples, clean small RNA reads were mapped to the wheat genome IWGSC RefSeq v1.0 [79] and *PGT* 21-0 genome [24] using bowtie 1.1.2 (alignment settings: no mismatches allowed –v0; report all alignments: -a –best –strata). To annotate high-confidence *Pgt* and wheat sRNAs from the sequencing data, we used the ShortStack software [80]. ShortStack predicts and quantifies sRNA-producing loci in a genome based on clusters of sRNA reads and miRNA-producing loci according to a series of tests, such as strandedness of the locus and the predicted precursor secondary structure.

### *Pgt* sRNA prediction, differential expression analysis and allelic sRNA prediction

To annotate and quantify high-confidence *Pgt* and wheat small RNAs from the sequencing data, we used the ShortStack 3.8.5 software [80] on the clean sRNA reads (--bowtie_m all). We further filtered the predicted sRNA clusters to include only those where >= 80% of reads are within 20-24 nts of length (recommended procedure in ShortStack to avoid degradation products) and where the cluster has >= 5 reads per million. The ShortStack software outputs sRNA cluster properties such as the most abundant sRNA (termed sRNA candidate) in the cluster, strandedness of the locus, miRNA annotation and phasing [80]. Strandedness of sRNA loci is determined by forcing the bowtie aligner to select one strand or the other with a probability that is proportional to the number of best sites on the strand. Stranded loci are typical of miRNA production in plants and are a requirement for annotation of a locus as a miRNA by ShortStack. We used the read counts returned by ShortStack for all predicted sRNA clusters and used edgeR [81] to assess which are differentially expressed at any of the infection stages versus germinated spores (FDR < 0.05, fold change > 2).

To assess if sRNAs have a homologous counterpart, we re-mapped the sequencing reads that define a sRNA locus to the remainder of the genome using bowtie 1.1.2 (alignment settings: two mismatches allowed –v2; report all alignments: -a –best –strata). If more than 25% of bases in a sRNA locus are covered by those mapped reads (using bedtools coverage version 2.28.0), it is marked as a candidate homolog. The sRNA locus with the highest coverage amongst the candidate homologs is returned as the predicted allelic counterpart. Circos 0.69.5 [82] was used to plot the links between homologous sRNAs across the chromosomes.

To assess the relationships of sRNAs and TEs, we re-mapped sRNAs to the genome using bowtie 1.1.2 (alignment settings: no mismatches allowed –v0; report all alignments: -a –best –strata). We reported repeats that overlap with those mapped sRNAs using bedtools intersect [83]. We then retrieved the genes that overlap with repeats using bedtools closest.

All plots were produced using Ggplot2 (Wickham, <u>2009</u>) and statistical significance was assessed with *t*□tests using the ggsignif package (https://cran.r-project.org/web/packages/ggsignif/index.html). Significance thresholds according to *t*□test are: NS, not significant; *, < 0.05; **, < 0.01; ***, < 0.001.

### Phylogenetic tree of RNAi genes

Argonaute protein sequences were aligned with MUSCLE 3.8.31 [84] and default parameters. FastTree 2.1.9 [85] was used to construct a phylogenetic tree from the protein sequence alignment (-pseudo -spr 4 -mlacc 2 -slownni). ETE 3.1.1 was used to draw the phylogenetic tree [86].

## Declarations

### Ethics approval and consent to participate

Not applicable

### Consent for publication

Not applicable

### Availability of data and materials

All scripts as well as code for generating the figures of this paper are available at https://github.com/JanaSperschneider/Publications_Code/tree/master/2019_12_StemRust_smRNA_Paper. Sequence data for the *Pgt* infection RNAseq is available at the National Center for Biotechnology Information Sequencing Read Archive under Bioproject PRJNA415866. Sequence data for the *Pgt* small RNAseq is available at CSIRO Data Access Portal under https://doi.org/10.25919/5bd93643b1e41. Hi-C data is available in NCBI under BioProject PRJNA516922. Methylation data is available at SRA under SUB7741177.

### Authors’ contributions

JS, JMT and PND conceived the study and designed experiments. JS analysed and interpreted all data sets and wrote the manuscript. AWJ, JN and BS performed methylation sequencing experiments and analysed methylation data. BX performed small RNA sequencing experiments. SJ, NMU and RM performed experiments and all authors contributed to data analysis and manuscript writing. All authors read and approved the final manuscript.

## Acknowledgements

We thank Xiaodi Xia for excellent technical assistance.

## Supplementary Figures

**Figure S1:**
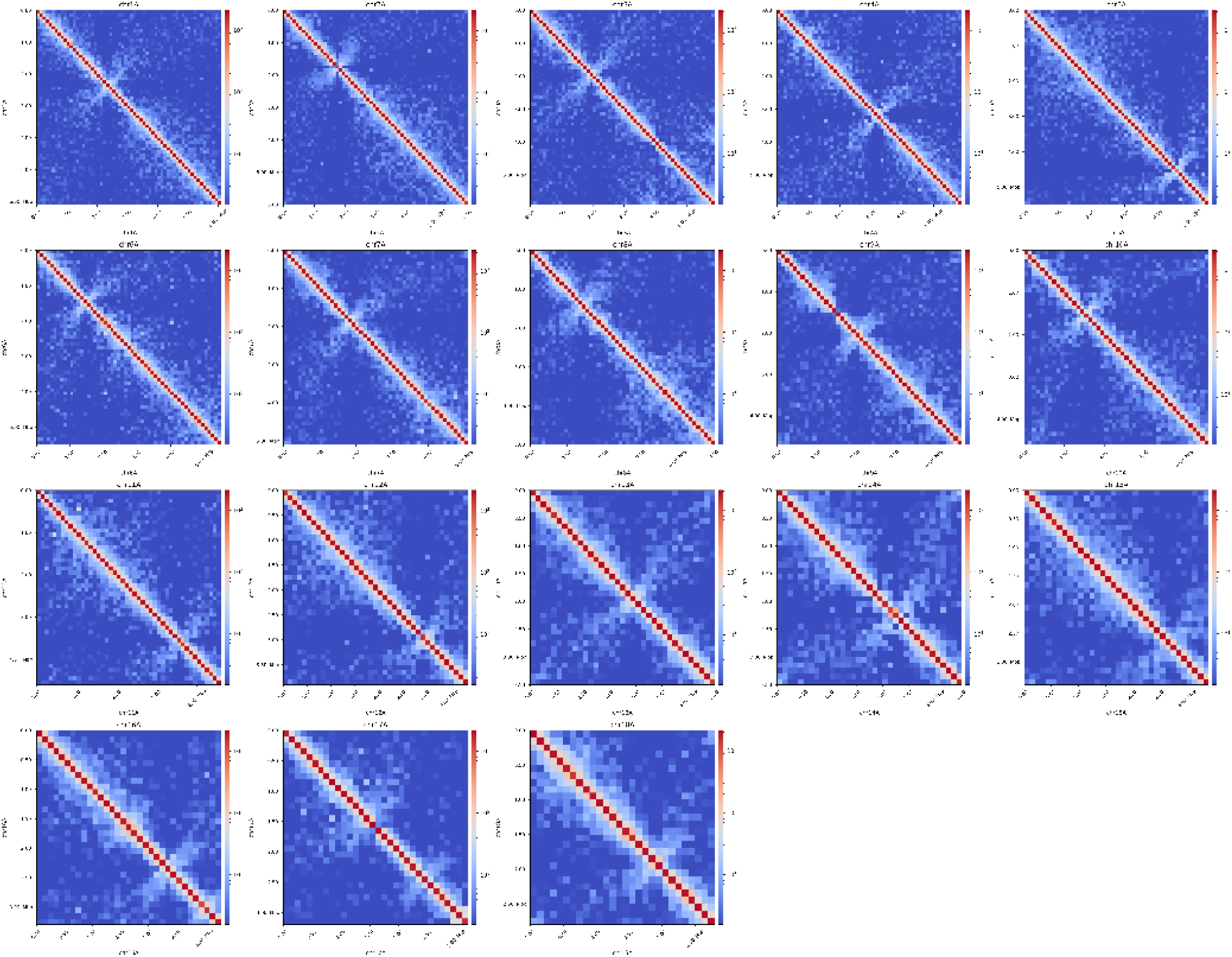
Hi-C contact maps for the 18 chromosomes of haplotype A show the presence of centromeres in each chromosome.

**Figure S2:**
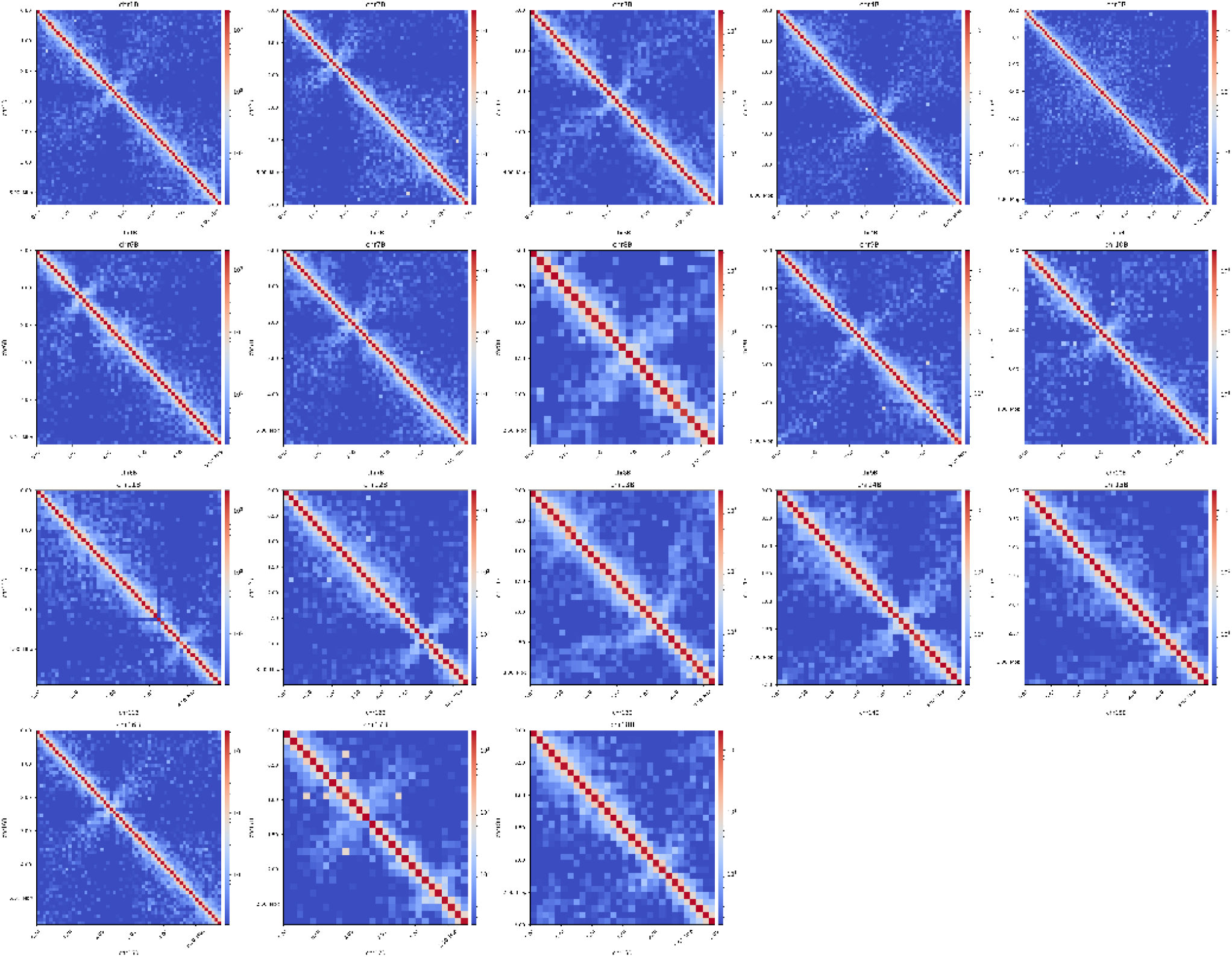
Hi-C contact maps for the 18 chromosomes of haplotype B show the presence of centromeres in each chromosome.

**Figure S3:**
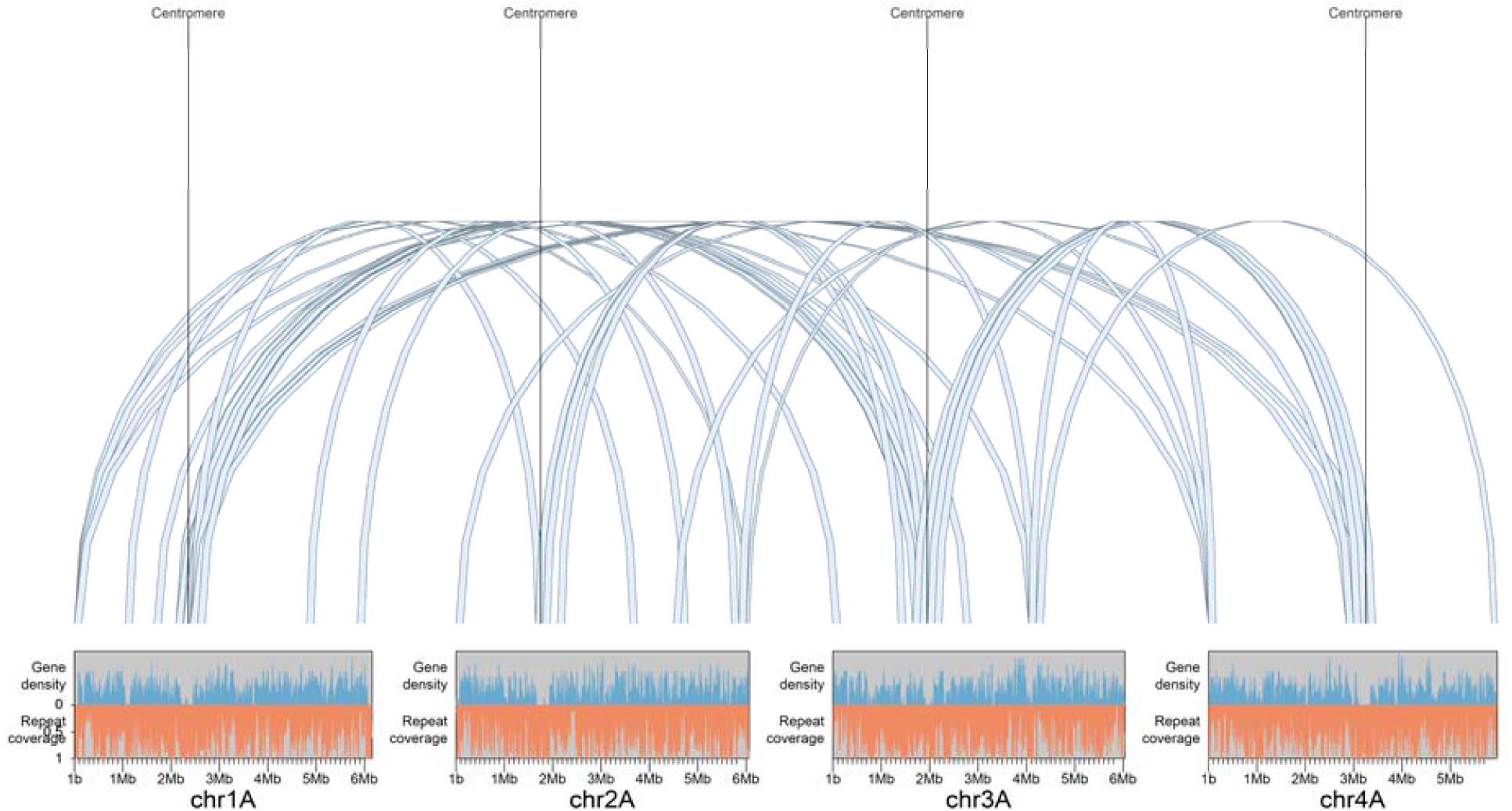
150 kbp bins with interaction frequency > 5 in the Hi-C interaction matrix are shown between chromosomes 1A, 2A, 3A and 4A. The putative centromeric regions share strong connections with each other. Densities of expressed genes and coverage of repetitive elements are shown with window size 10 kbp. The centromeric regions are gene-poor regions with high repetitive element coverage.

**Figure S4:**
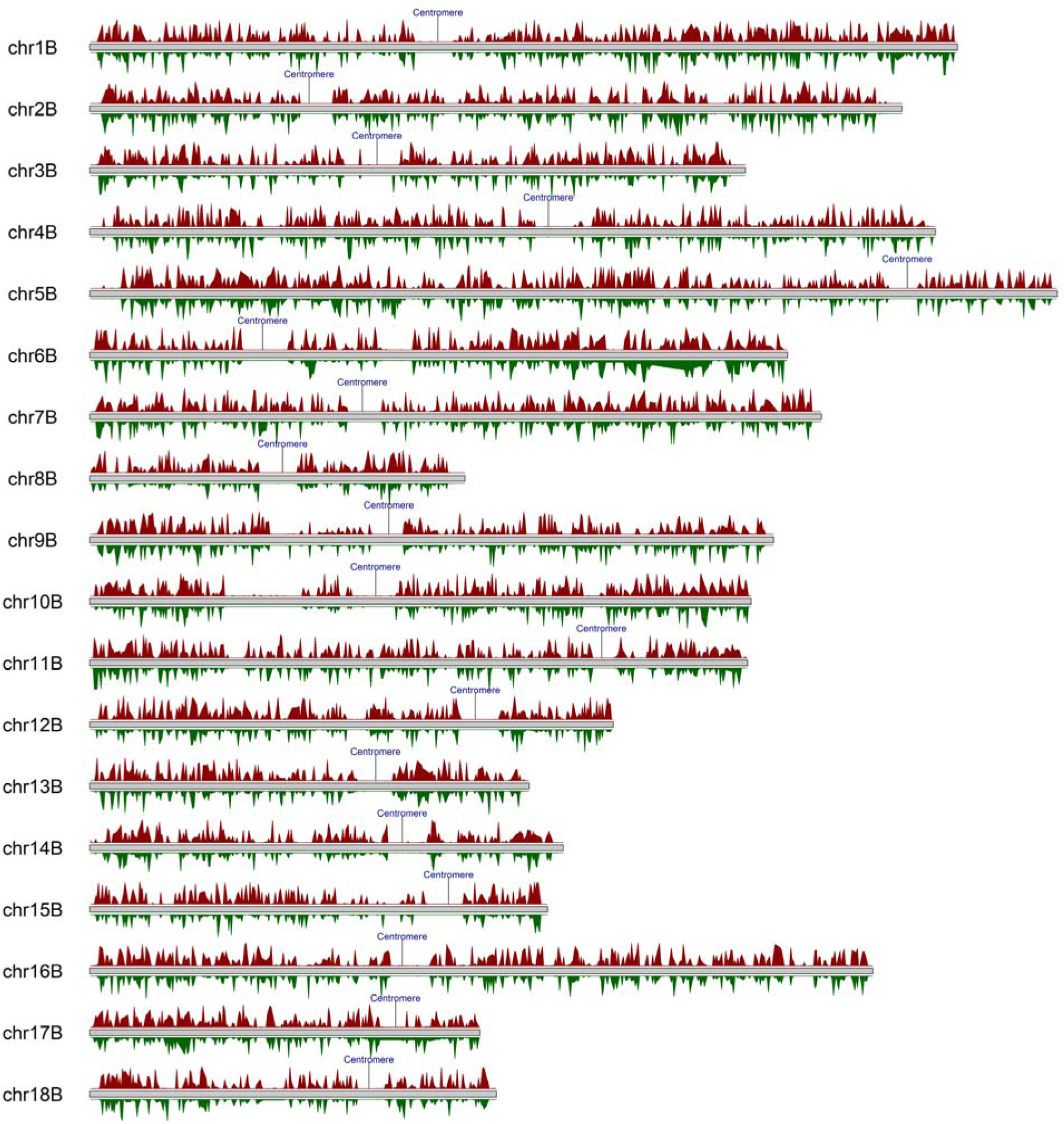
The positions of the centromeres in haplotype B as indicated by the Hi-C contact map are in transcriptionally silent genomic regions. Reads per million (RPM) for the late infection (7 dpi) and germinated spores RNAseq samples are shown in red and green, respectively (10 kb windows, RPM from 0-100 are shown for clarity).

**Figure S5:**
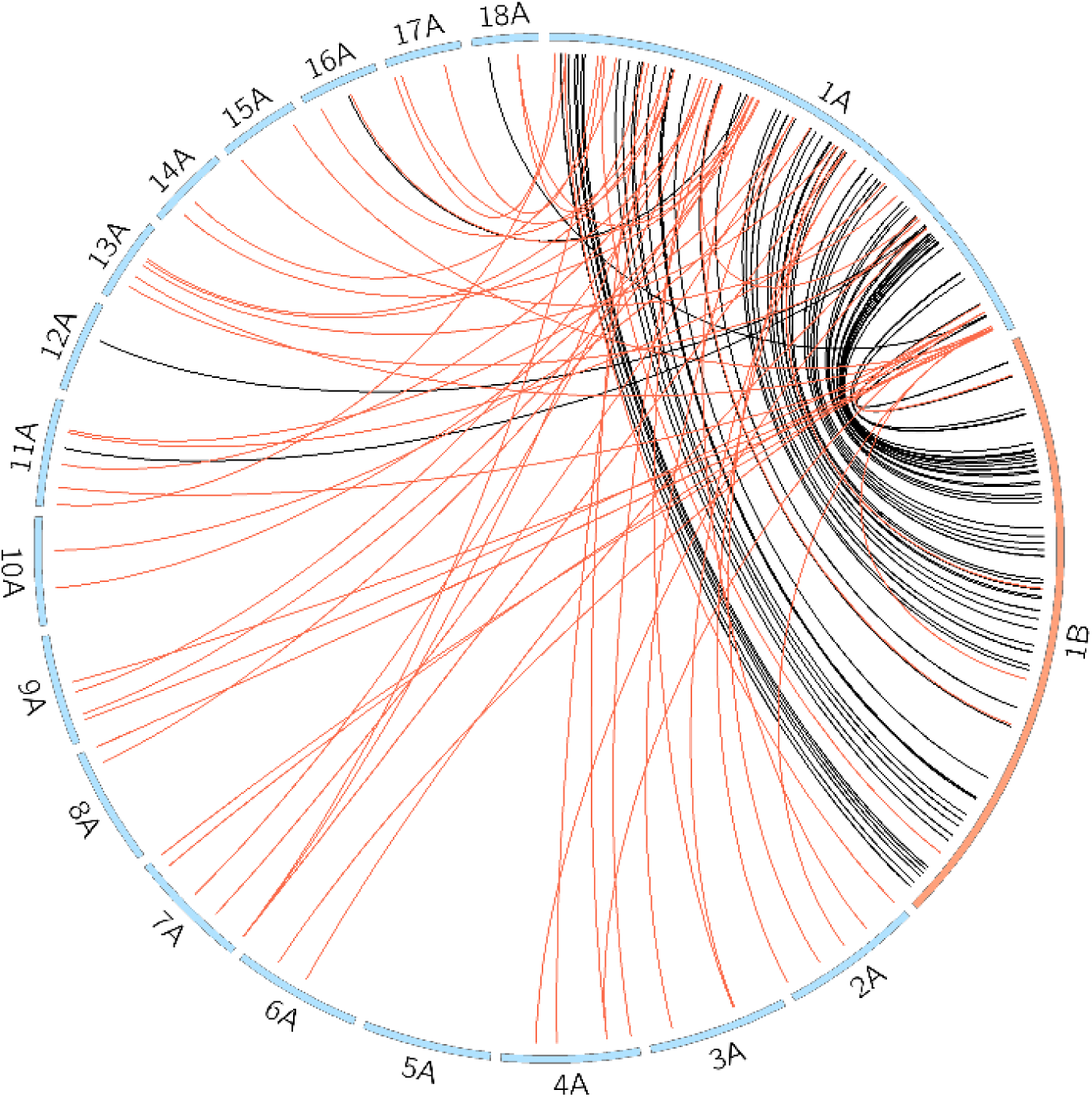
*Pgt* allelic sRNA pairs and their genomic localization for chromosome 1A. *Pgt* sRNAs that are up-regulated in germinated spores (late infection) and their homologous counterparts are shown with black (red) links. sRNAs that are up-regulated in germinated spores appear to be in syntenic on the two haplotype chromosomes 1A and 1B (shown at twice their size, other chromosomes shown at 0.2 their size). In contrast, sRNAs that are up-regulated during late infection on chromosome 1A have homologous counterparts on all other chromosomes except 5A and 12A.

## Supplementary Files

Supplementary File S1: FASTA file of predicted *Pgt* siRNAs.

Supplementary File S2: FASTA file of predicted *Pgt* miRNAs.

Supplementary File S3: FASTA file of predicted wheat siRNAs.

Supplementary File S4: FASTA file of predicted wheat miRNAs.

Supplementary File S5: FASTA file of *Pgt* sRNAs predicted to be up-regulated in germinated spores.

Supplementary File S6: FASTA file of *Pgt* sRNAs predicted to be up-regulated in 3 dpi and/or 5 dpi.

Supplementary File S7: FASTA file of *Pgt* sRNAs predicted to be up-regulated in 7 dpi.

Supplementary File S8: FASTA file of *Pgt* sRNAs predicted to have no differential expression.

